# Automated Parameter Estimation for Camera Trap Density Models Using Computer Vision-Enhanced Distance Sampling

**DOI:** 10.64898/2026.06.14.732225

**Authors:** Sierra McMurry, Ben Goldstein, Mohammed Alyetama, Roland Kays

## Abstract

Models for estimating animal density from camera traps require four parameters informing detection: movement speed, daily activity level, staying time (duration animals remain within the detection zone), and effective detection distance. These parameters traditionally come from labor-intensive manual measurements and auxiliary telemetry. Recent advances in computer vision can provide the positions of animals in camera trap images, which have been used for distance sampling. We extend this approach to extract all four parameters from imagery, providing the first AI-derived estimates of movement speed and staying time from automated coordinate tracking. We also introduce a new joint multi-species hierarchical distance function that estimates deployment-specific effective detection distances while borrowing strength across species through partial pooling.

Our pipeline integrates MegaDetector for animal detection, the Segment Anything Model for segmentation, and Dense Prediction Transformers for monocular depth estimation. From frame-level coordinates, we reconstruct movement trajectories across burst sequences to estimate speed with size-biased distribution corrections, calculate staying time through bounding box interpolation, and estimate activity levels from detection timestamps. The joint hierarchical distance function decomposes the detection scale parameter into a shared deployment-level effect and species-specific offsets, so species effects represent deviations from the multi-species average, allowing data-rich species to inform detection conditions where rare species have few observations. AI-derived scene depth enters the model as a covariate on detection range, providing a vegetation openness metric from the same pipeline. To address position errors from depth estimation, we apply data quality filters.

We processed 122,574 frames from 181 deployments across montane forests in Washington and Montana, generating parameter estimates for 12 species without manual annotation. Automated speed estimates produced day ranges 2.7 to 4.3 times GPS telemetry-derived daily distances, reflecting differences between encounter velocity within detection zones and landscape-scale displacement. Deployment-level variation in detectability exceeded species-level differences 3:1, with scene depth strongly predicting detection range; mean effective detection distances ranged from 4.1 to 7.6 m. Applied to a Random Encounter Model, these parameters yielded a white-tailed deer density estimate of 21.4 animals/km² and the Random Encounter Staying Time model yielded 11.6animals/km² in Montana. This pipeline enables scalable density estimation across large camera trap networks.

## I. Introduction

Camera trap arrays have transformed wildlife monitoring, enabling simultaneous data collection on mammal communities. Through standardized programs, these arrays are now deployed repeatedly across cities (Magle et al. 2019), countries (Rooney et al. 2025), and continents (Kays et al. In Review), generating annual datasets of unprecedented spatial and temporal scope. These data enable powerful comparisons across space and time while capturing community-level dynamics through passive, non-invasive sampling. However, most analyses from these programs have relied on occupancy or relative abundance indices rather than density, limiting the strength of inference that can be drawn about population status.

Population density represents the gold standard metric for wildlife monitoring, enabling direct comparisons across space and time and robust inference about conservation status unlike occupancy or relative abundance indices, which yield only relative measures (Callaghan et al., 2024). Recent advances in unmarked density models have made density estimation possible from camera trap data without individual identification (Moeller et al. 2018, Gilbert et al. 2021, Palencia et al. 2021, Loonam et al. 2021). However, despite these advances, practical implementation remains severely limited by labor-intensive requirements for parameter measurement.

Beyond just counting animal detections, all camera trap density models require additional parameters that must be measured for accurate estimation. First, all models require measurement of the effective area surveyed by each camera. Detection distance was initially estimated using human-led field trials (Rowcliffe et al., 2008) where a person approached the camera at different speeds and directions. Rowcliffe et al. (2011) showed that animal positions could be measured in the field by using local landmarks in photos as reference points, enabling a distance sampling approach to estimate detection distance while accounting species-specific detection differences, for example, the strong effect of larger animals being detected further away. Various approaches have been suggested for making distance measurements more efficient, including placing measuring sticks at known distances (Howe et al., 2017) or noting whether animals pass specific markers (Haucke et al., 2022, Palencia et al., 2021), but none have been widely adopted due to the extra work involved and concerns about markers affecting animal behavior (Becker et al., 2022). More recently, viewshed-based approaches have also been developed to estimate the viewable area for each camera based on field conditions, defining the sampling area as a circular sector determined by the camera’s lens angle and maximum visible distance identified from field landmarks (Moeller et al., 2018). However, the viewshed approaches measure the maximum visible area for a camera without accounting for the actual detection zone, which could be shorter depending on the motion sensor’s ability to detect moving animals, which can vary depending on animal size, ambient temperature, and movement speed.

Additionally, some density models require measures of animal movement speeds (Rowcliffe et al. 2008). These are typically drawn from GPS tracking studies, although rarely from the same population being camera trapped (Palencia et al. 2022). Rowcliffe et al. (2016) suggested that a population average movement speed could be derived from camera trap data by measuring instantaneous speeds of animals moving through the camera trap view and then adjusting this by the percent of the day the animal was active which can also be derived from the temporal distribution of camera detections (Rowcliffe et al. 2014). Traditionally these movement parameters had to be acquired through manual measurements that are time-consuming and limit both the efficiency and scalability of multi-species density studies. Thus, camera trap images themselves could theoretically provide all the data needed to estimate density from unmarked animals — detection rates, staying time, movement speeds, and area sampled — but with substantial extra work.

Recent advances in artificial intelligence present unprecedented opportunities to automate these measurements, potentially achieving orders of magnitude improvements in processing efficiency. Computer vision (CV) tools that record the position of an animal in an image (Haucke et al. 2022) suggest a new potential to generate these data automatically, Henrich et al. (2024) used these in a semi-automated distance sampling density estimate and others reduced the calibration burden for distance estimation (Henrich et al. 2024, Aamir et al. 2025).However, these applications have focused on distance estimation alone, leaving movement speed, staying time, and activity level still dependent on manual measurement or auxiliary telemetry. Even within distance estimation, several technical challenges remain unresolved. These challenges include depth estimation errors in dense vegetation, where monocular models rely on learned spatial priors that tend to over-smooth fine-scale depth structure (Haucke et al. 2022); loss of tracking accuracy under partial occlusion or limited ground context, which is well documented in computer vision tracking benchmarks (Smeulders et al. 2014) and likely exacerbated in camera trap imagery and temporal instability between frames that can compromise individual tracking. Collectively, these limitations have prevented the full automation of density estimation from camera trap imagery.

This paper addresses these limitations by presenting an integrated pipeline that uses CV to measure all four essential parameters for camera trap density estimation: effective detection distance, staying time, movement speed, and activity level. Building on recent advances in automated distance estimation (Haucke et al. 2022), we implement automated sign detection for deployment calibration, precise animal segmentation using the Segment Anything Model (SAM) (Kirillov et al. 2023), and Dense Prediction Transformer (DPT)-based depth estimation (Ranftl et al. 2021) for both distance measurement and frame-level coordinate extraction. Because monocular depth estimation introduces position errors that can propagate through trajectory and distance calculations, we develop a series of quality filters — including burst-frame collapse, coordinate anchoring, edge-frame removal, and tortuosity thresholds — to remove obviously poor position estimates while retaining the majority of observations. From these coordinates, we reconstruct movement trajectories to estimate speed using the size-biased distribution correction (Rowcliffe et al. 2016) and calculate staying time through bounding box interpolation. We also make full use of this large dataset of animal positions to estimate deployment-specific effective detection distances via a new joint multi-species hierarchical detection model that borrows strength across species within deployments. This is a powerful new extension of the distance sampling framework of Howe et al. (2017) that takes advantage of the multi-species CV-derived depth estimates to estimate effective detection distance for each species at each camera deployment. We demonstrate this pipeline across camera deployments in Washington and Montana, generating automated parameter estimates for 11 mammal and one avian species. Our pipeline advances the field in three ways. First, it is the first to automate all four density parameters (EDD, speed, activity, staying time) from camera trap imagery, whereas previous work automated only distance estimation (Haucke et al. 2022, Henrich et al. 2024, Aamir et al. 2025). Second, the joint multi-species hierarchical detection model provides deployment-specific EDD estimates that account for site-level variation in detection conditions, a capability not available from conventional single-species distance sampling. Third, by eliminating manual measurement requirements, the pipeline enables community-level density estimation at scales previously impractical.

## II. Methods

### 2.1 Study Design

#### 2.1.1 Study Area

Camera trap surveys were conducted across deployment locations in montane forest ecosystems of Washington and Montana, USA (approximately 47°–49°N, 113°–124°W). The Washington study sites included locations within the Olympic Peninsula, North Cascades, and eastern Cascade forests.

Vegetation communities consisted primarily of mixed conifer stands typical of the inland Northwest and Cascade Range. Forest composition varied with elevation and aspect, transitioning from Douglas-fir (*Pseudotsuga menziesii*) and ponderosa pine (*Pinus ponderosa)* associations at lower elevations to subalpine fir and mountain hemlock at higher sites (Franklin & Dyrness 1988). Understory vegetation included huckleberry, rhododendron, and devil’s club in mesic sites, with bunchgrasses and shrub-steppe species in drier locations. Survey locations ranged from 300 to 1,800 meters in elevation, capturing the full environmental gradient used by the regional mammal assemblage. The study areas support populations of white-tailed deer (*Odocoileus virginianus*), mule deer (*O. hemionus*), elk (*Cervus canadensis*), moose (*Alces alces*), American black bear (*Ursus americanus*), mountain lion (*Puma concolor*), coyote (*Canis latrans*), bobcat (*Lynx rufus*), and various smaller mammals.

**Figure 1.**
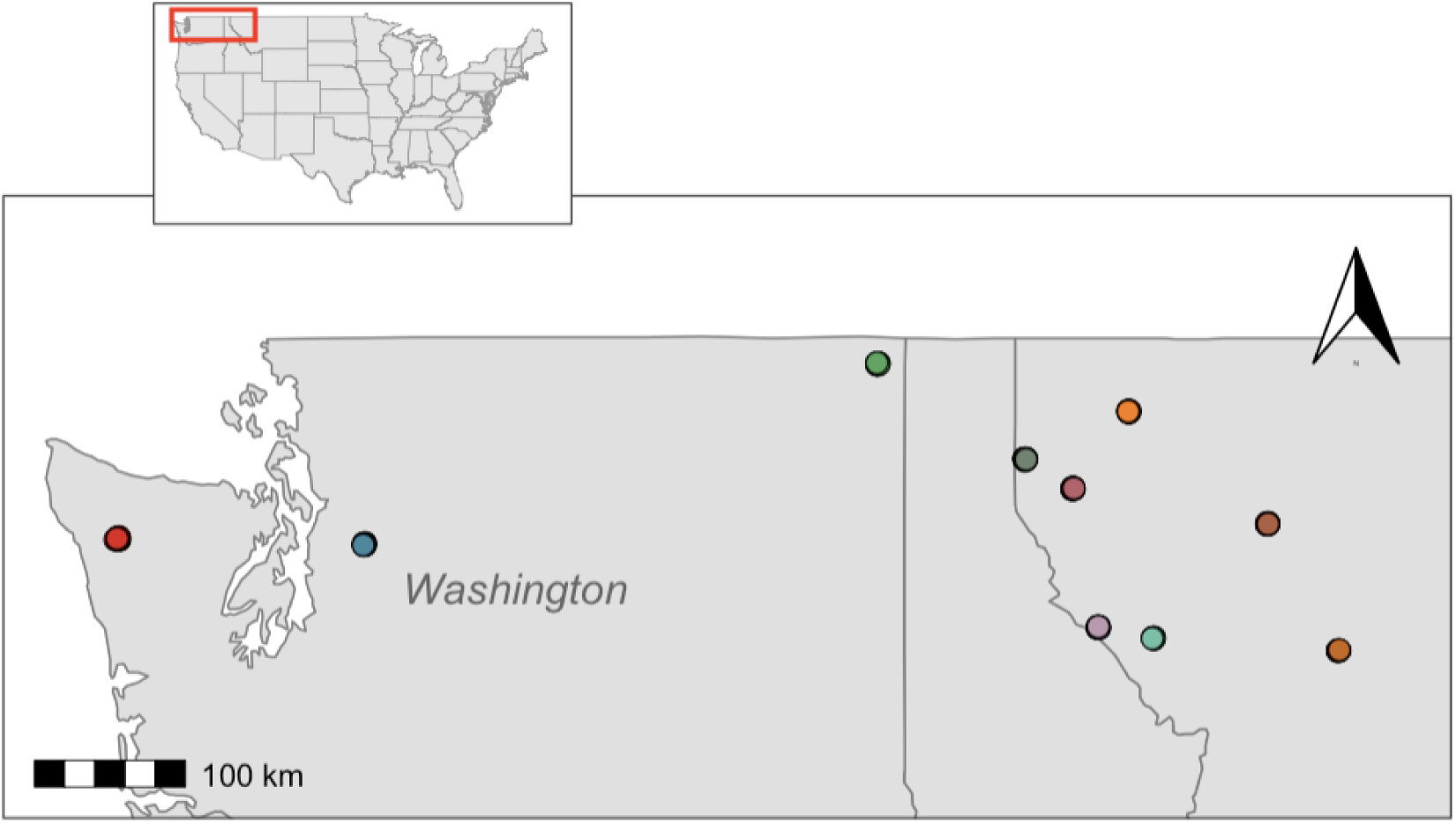
Study area map showing 10 camera trap arrays in Washington and Montana, USA. Arrays are colored individually on the map. Inset shows study area location within the contiguous United States.

#### 2.1.2 Camera Deployment Strategy

We deployed cameras on U.S. Forest Service lands using stratified site selection based on forest type. Within each selected forest type, we chose a site location at random, and placed cameras in a systematic grid of 30 units spaced at 300 m intervals to ensure consistent spatial coverage and maximize detection probability. We did not use attractants or bait to avoid biasing animal behavior or movement patterns. Each deployment consisted of a single camera mounted at approximately 1 meter height on trees oriented parallel to the ground. We utilized three camera models: Browning Recon Force Elite HP5, Reconyx Hyperfire 2, and Reconyx HC500 Hyperfire. Cameras operated continuously in motion-trigger mode with manufacturer-default sensitivity settings. Each deployment was maintained for a minimum of 30 days.

#### 2.1.3 Field Calibration Protocol

While AI models can automatically estimate relative position from an image, calibration of each camera deployment is needed to enable it to report location in real distance units. Different camera models have distinct optical characteristics that directly affect depth estimation accuracy, requiring model-specific parameters for accurate calibration (Appendix S1: Table 1). Our approach follows the general strategy of placing reference objects at known distances (Rowcliffe et al. 2011, Howe et al. 2017, Haucke et al. 2022), but automates sign detection to eliminate manual annotation of reference images.Our field calibration (Box 1) uses 30 × 30 cm white signs at 3 m intervals, extending one interval beyond maximum trigger distance.

**Table 1.** Summary of quality filters applied to distance estimation (frame-level) and movement speed estimation (sequence-level). Percentages are relative to the total at each level. Bbox >80% of frame is the AI-detected bounding box that occupies more than 80% of the frame area, indicating the animal is too close to the camera for reliable depth estimation. Too close (world_y < 1 m) is the AI-estimated forward distance to the animal is less than 1 m, outside the reliable range of the depth model. Too far (world_y > 26m) is the AI-estimated forward distance exceeding 26m, the maximum distance supported by our depth calibration data beyond which depth estimates become unreliable. Frame speed (>20 m/s) is the displacement between consecutive frames divided by elapsed time exceeding 20 m/s, used as a threshold to flag coordinate errors rather than biologically real movement.

| Filter | Criterion | n Removed | % |
| --- | --- | --- | --- |
| **Frame Level** |  |  |  |
| Bbox >80% of frame | bbox_area_pct > 80% | 2,896 | 2.5% |
| Too close | world_y < 1 m | 1,963 | 1.7% |
| Too far | world_y > 26 m | 4,262 | 3.6% |
| Impossible speed | frame_speed > 20 m/s | 444 | 0.4% |
| Total frames flagged | Any of above | 8,845 | 7.5% |
| **Sequence Level** |  |  |  |
| Bbox sequence excl. | Any frame bbox >80% | 281 | 7.8% |
| Group size >1 | group_size > 1 | 995 | 27.6% |
| Zero duration | duration = 0 s | 568 | 20.9% |
| Tortuosity exceeded | Species-specific threshold | 388 | 14.3% |
| Zero/invalid speed | speed = 0 or NA | 800 | 29.5% |
| Retained for SBD | Passed all filters | 1,524 | 56.2% |

### 2.2 AI Pipeline

We created an analytical workflow for calibrating the scene for each camera, detecting and segmenting the animals in each image, and estimating their position relative to the camera (Figure 2).

**Figure 2.**
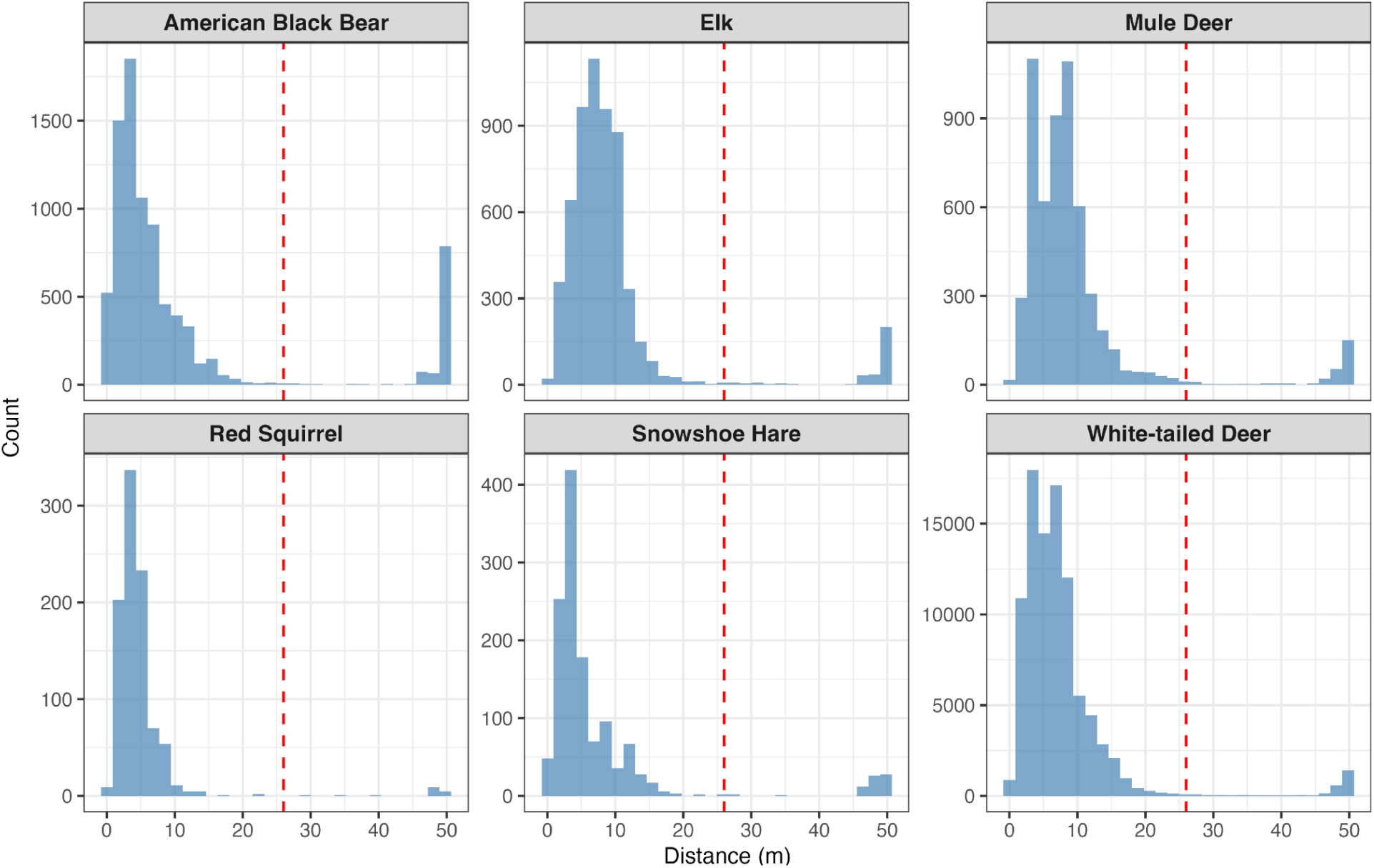
Distance Distribution by Species. The bright red line is 26 m (our distance cut off). Detections at 50 m are often from animals close to the camera lens and the AI chooses their position at the maximum distance. All distance estimates further than 26m are disregarded.

#### 2.2.1 Sign Detection and Depth Calibration

Accurate distance estimation requires deployment-specific calibration to convert relative depth values from the AI model into real-world distances. Previous approaches relied on either permanent field markers with manual image comparison or physical return visits to measure detection distances (Haucke et al. 2022), both of which are labor-intensive and limit scalability. We automated this step using a custom detection model (SignD) (Alyetama 2025) that automatically locates the white square calibration signs within reference images and reads the distance number displayed on each sign. The SignD model generates precise bounding boxes around detected signs and creates corresponding binary masks utilized in subsequent depth calibration steps. This model has demonstrated high accuracy in sign detection across our dataset, successfully handling variations in sign presentation and lighting conditions.

The depth calibration procedure employs a pre-trained Dense Prediction Transformer (DPT) model (Haucke et al. 2022) for initial monocular depth estimation on the reference images. For each camera deployment, our automated calibration application built using Prefect requires only the input CSV file and the corresponding camera specifications from Table 1 (Table S1) to complete the entire calibration process with minimal manual intervention. The system automatically extracts and applies model-specific parameters including horizontal and vertical field of view, focal length, and sensor specifications, incorporating these into deployment-specific configuration files with additional processing thresholds (minimum detection distance of 0.5 meters, maximum reliable detection range of up to 26 meters).

#### 2.2.2 Animal Detection, Segmentation, and Position Estimation

The distance estimation process applies the DPT depth estimation model (Haucke et al. 2022) to generate a depth map of each image, calibrated using reference depth maps established during the calibration workflow. Monocular depth estimation for camera trap applications has advanced rapidly; Niccoli et al. (2025) recently introduced the first benchmark dedicated to wildlife monitoring conditions, evaluating four state-of-the-art models on camera trap imagery with ground truth distances from calibrated ChARUCO patterns. Their results demonstrated that newer architectures such as Depth Anything V2 can achieve substantially lower error (MAE 0.454 m) than earlier approaches, while also revealing that models optimized for urban or indoor scenes may degrade significantly in natural outdoor environments. Our pipeline uses the DPT architecture (Ranftl et al. 2021) paired with deployment-specific calibration, which mitigates the domain gap that Niccoli et al. identified as a fundamental challenge for zero-shot MDE models in wildlife settings.

A key challenge is precisely and consistently determining an animal’s position within this frame. We use MegaDetector (Beery et al. 2019) to identify where animals are within the frame and create bounding boxes around each individual. Within these bounding boxes, we then apply the Segment Anything Model (SAM) (Kirillov et al. 2023) to precisely segment the animal’s shape, allowing us to isolate pixels corresponding to the animal itself from the background within the bounding box. This segmentation step goes beyond the bounding-box-only approaches evaluated in previous work (Haucke et al. 2022; Niccoli et al. 2025), reducing contamination from background pixels that are inevitably included in the rectangular bounding box region.

The final challenge is to consistently measure the distance to the same part of the segmented animal. To address this, we extract depth values from all pixels within the SAM-segmented animal region and calculate the 20th percentile of this distribution. Within any segmented region, depth values comprise a mixture of pixels on the animal itself (at the true distance) and background pixels visible through imperfect segmentation boundaries (at greater distances). Taking the mean of all pixels biases the estimate toward the background, while very low percentiles (e.g., 5th) risk capturing foreground noise or segmentation artifacts. The 20th percentile focuses on the nearest cluster of pixels, which corresponds to the animal’s body, while remaining robust to occasional outliers. This method follows Haucke et al. (2022), who demonstrated that percentile-based approaches outperform mean or median methods for bounding box distance estimation. Niccoli et al. (2025) similarly found that median-based depth extraction consistently outperformed mean-based approaches across all models tested, supporting the general principle that central or lower-quantile extraction reduces sensitivity to background contamination and outlier depth values.

Beyond distance estimation, our pipeline extends the Haucke et al. (2022) framework by extracting two-dimensional position coordinates (lateral position and depth) for each detection, rather than a single scalar distance. This enables reconstruction of movement trajectories across sequential frames within burst sequences, providing the positional data required for speed estimation (Section 2.3) and staying time calculation (Section 2.5). The improved accuracy demonstrated by newer depth models (Niccoli et al. 2025) suggests that future implementations could achieve better coordinate precision by adopting architectures such as Depth Anything V2 while retaining the same analytical framework for parameter estimation.

#### 2.2.3 Quality Filtering and Edge Case Handling

By comparing the estimated position with the original photographs we found some situations where the depth estimation consistently performed poorly. Therefore we implemented filtering criteria separately for distance estimation and movement speed calculation, prioritizing data retention while ensuring biological plausibility.

We flag and filter detections in two scenarios where depth estimation becomes unreliable:

1. **Bounding box size** — Detections where the bounding box occupies >80% of frame area indicate the animal is too close for reliable depth estimation. At this proximity, limited background context causes erratic estimates such as an animal at 1m registering as max distance.
2. **Maximum distance** — Detections beyond 26 m exceed our maximum reliably calibrated depth range and are presumed to be erroneous estimates and are excluded.

Combined, these filters affected approximately 7.5% of frame-level detections. Removing flagged detections changed overall effective detection distance (EDD) by <1%, confirming that exclusion does not introduce systematic bias in distance estimation.

##### Movement Speed Filters

For speed estimation, we apply additional filters at the sequence level, some following Rowcliffe et al. (2016):

1. **Multi-frame requirement** — Sequences must contain ≥2 frames for trajectory calculation. Single-frame detections were excluded; manual inspection confirmed these primarily resulted from cameras failing to capture multiple frames of distant, moving animals rather than representing a distinct behavioral class.
2. **Biological plausibility** — Sequences with speeds >20 m/s were removed as physically impossible for terrestrial mammals (Garland 1983).
3. **Sequence-level bounding box exclusion** — If any frame within a sequence contained a bounding box exceeding 80% of frame area, the entire sequence was excluded from speed analysis. Even a single unreliable coordinate within a sequence contaminates the path distance calculation, so sequence-level exclusion is necessary despite the frame-level filter above.
4. **Single-animal requirement** — Sequences where group size exceeded one were excluded from speed estimation. When multiple animals are present, AI-tracked coordinates can jump between individuals across timestamps, producing path distances that reflect inter-animal spacing rather than individual movement. At the camera’s low frame rate (∼1.4 fps), reliable individual tracking within groups is not possible. Future work with higher frame rate video could overcome this limitation.

**Figure 3.**
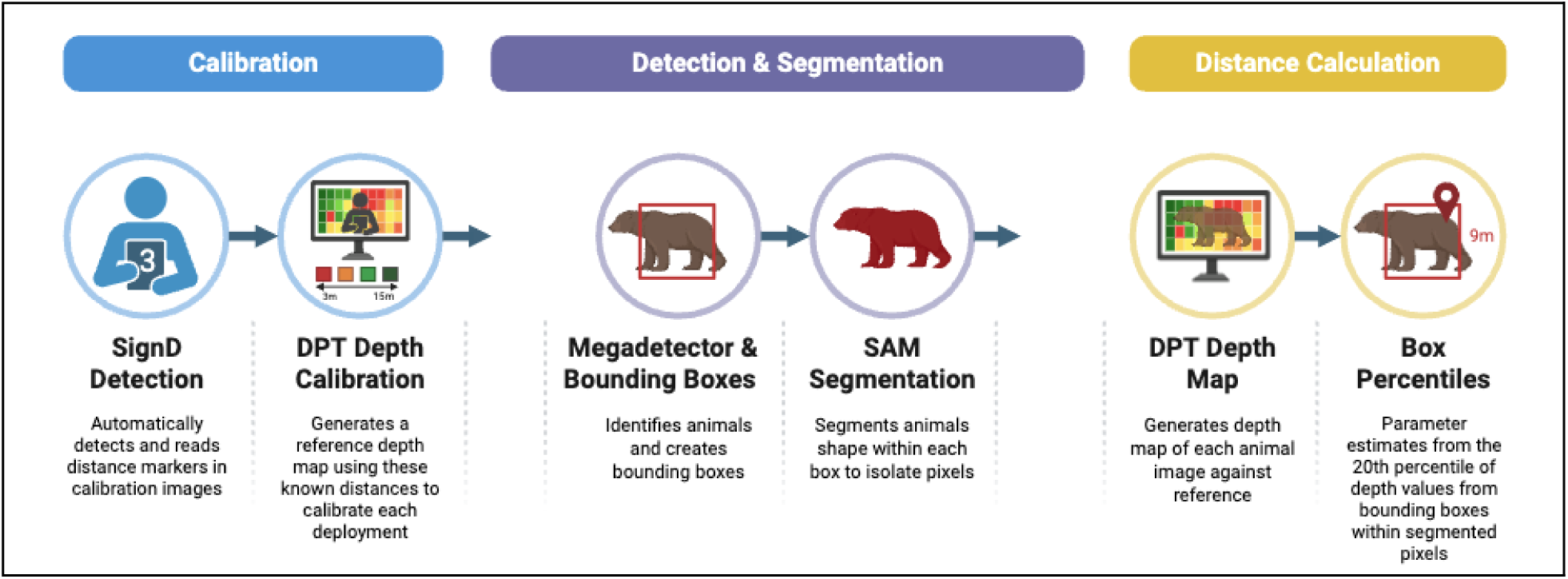
AI Distance Pipeline Workflow. Schematic of the automated pipeline for calibration, detection, segmentation, and distance estimation from camera trap imagery. Calibration signs placed at known distances are detected by the SignD model and used to create deployment-specific depth references. For wildlife images, MegaDetector identifies animals and produces bounding boxes, SAM segments the animal from background, and the DPT depth model estimates distance using the 20th percentile of depth values within the segmented region. Outputs include world coordinates (world_x, world_y) for each detection in each frame.

### 2.3 Movement Speed Estimation

#### 2.3.1 Coordinate Extraction and Trajectory Reconstruction

The AI pipeline generates world coordinates (world_x, world_y, in meters) relative to the camera trap for each detected animal in each frame, based on the estimated position from the segmented animal region (see Section 2.2.3). This approach enables automated extraction of movement trajectories from camera trap sequences.

*Burst frame corrections* — For multi-frame sequences, we reconstructed movement paths by connecting world coordinates across consecutive frames. Camera traps operating in burst mode capture multiple frames within the same second, all sharing an identical timestamp. Small AI coordinate shifts between successive burst images accumulate as artificial movement, inflating path distance even when the animal barely moved. To address this, we collapsed each sequence to one coordinate per unique timestamp, retaining only one frame per second. Path distance was then calculated between these second-level positions only, and sequence duration was computed as the time elapsed between the first and last unique timestamp. Speed could not be calculated for sequences with only one unique timestamp.

*Frame-edge AI errors* — When visually comparing position estimates to photos of the animals we noticed the AI position estimation was occasionally less accurate at the right or left edge of the image, indicating a large movement in the y direction when none was actually present. Therefore, we created a filter to remove points that were at the edge of the frame (i.e. bounding box extended into the outer 1/6 of the frame on the right or left side) and had a large movement in the y direction compared to the rest of the trajectory (i.e. depth movement exceeded 1.5× the median depth of non-edge frames in the same sequence). Edge frame movements with reasonable depths were retained.

*Point Anchoring* — The AI estimated positions of a stationary animal could still change slightly just due to changes in their posture (aka AI coordinate jitter). Therefore, we used point anchoring to ignore small movements until the estimated position moved beyond a predefined distance threshold, at which point the anchor location was updated. Because the magnitude of this coordinate noise scales with body size; larger animals produce larger bounding boxes and thus larger apparent displacements from posture changes alone, we used species specific thresholds.

Rather than arbitrarily assigning thresholds solely from body-size categories, we empirically determined optimal thresholds for each species using a sweep-and-elbow approach. For each species, we calculated size-biased distribution speed across a range of candidate thresholds (0.1 to 1.5 m) and identified the threshold at which the speed-vs-threshold curve stabilized using the Kneedle algorithm (Satopää et al. 2011), which detects the point of maximum curvature in the monotone decreasing portion of the curve. Below this threshold, raising the anchor reduces noise-driven speed inflation; above it, further increases remove real movement and either plateau or increase speed by selectively excluding slow sequences. We applied two reliability checks: if the curve was inverse (speed increased with threshold, indicating the threshold made estimates worse) or if fewer than 10 sequences survived at the detected elbow, the species was flagged and reverted to a body-size default.

*Path tortuosity filter* — When comparing the AI position estimates with the movement seen in sequences of images we still noticed occasional erroneous AI estimates suggest an animal rapidly away from the main movement path, and then returns, thus artificially inflating the movement distance. These create highly tortuous paths that are not typical of normal animal movements (e.g. Appendix S1: Figure S2). Therefore, we applied a species-specific tortuosity threshold to exclude sequences where accumulated coordinate noise above the anchoring threshold produced biologically implausible paths. Tortuosity was calculated as the ratio of path distance to straight-line distance (first frame to last frame). For most species, sequences exceeding a tortuosity ratio of 3 were excluded, as no animal traveling through a detection zone produces a path three times longer than its net displacement within a single short sequence. For small mammals (red squirrel, snowshoe hare), which exhibit genuinely erratic foraging and escape movements, the threshold was relaxed to 5. Elk were assigned a tortuosity threshold of 4 based on their movement characteristics.

*Straight-line distance* — After the above corrections and filters were applied we used frame-to-frame distances above each species’ threshold were calculated as Euclidean distances between consecutive coordinate pairs. The total path distance was computed as the sum of these segments, and the speed was this total path distance divided by the total time of the sequence.

#### 2.3.2 Size-Biased Distribution Framework

One fundamental challenge in camera-based speed estimation arises from detection bias inherent in camera trap sampling. As Rowcliffe et al. (2016) demonstrated, the probability of detecting an animal is proportional to its speed because faster-moving animals encounter cameras more frequently and are therefore overrepresented in the sample. This creates a ‘size-biased’ distribution where each animal’s probability of detection is proportional to its speed (the ‘size’ variable in this context). Under this size-biased sampling, the arithmetic mean of observed speeds overestimates the true population mean. To correct for this bias, Rowcliffe et al. (2016) showed that the harmonic mean of observed speeds provides an unbiased estimate of the population mean speed. The harmonic mean speed (H) was calculated as:

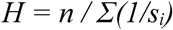

where n is the number of movement sequences and sᵢ is the speed of sequence i. This harmonic mean provides an unbiased estimate of population mean speed under size-biased sampling conditions (Rowcliffe et al. 2016).

The harmonic mean effectively down-weights higher values, counteracting the oversampling of faster speeds. This SBD (Size-Biased Distribution) framework assumes that detection probability scales linearly with speed, though in practice this relationship may be modified by sensor characteristics such as trigger speed, detection zone geometry, and recovery time between captures.

We calculated 95% confidence intervals using 1,000 bootstrap replicates for each species with ≥10 valid sequences.

**Figure 4.**
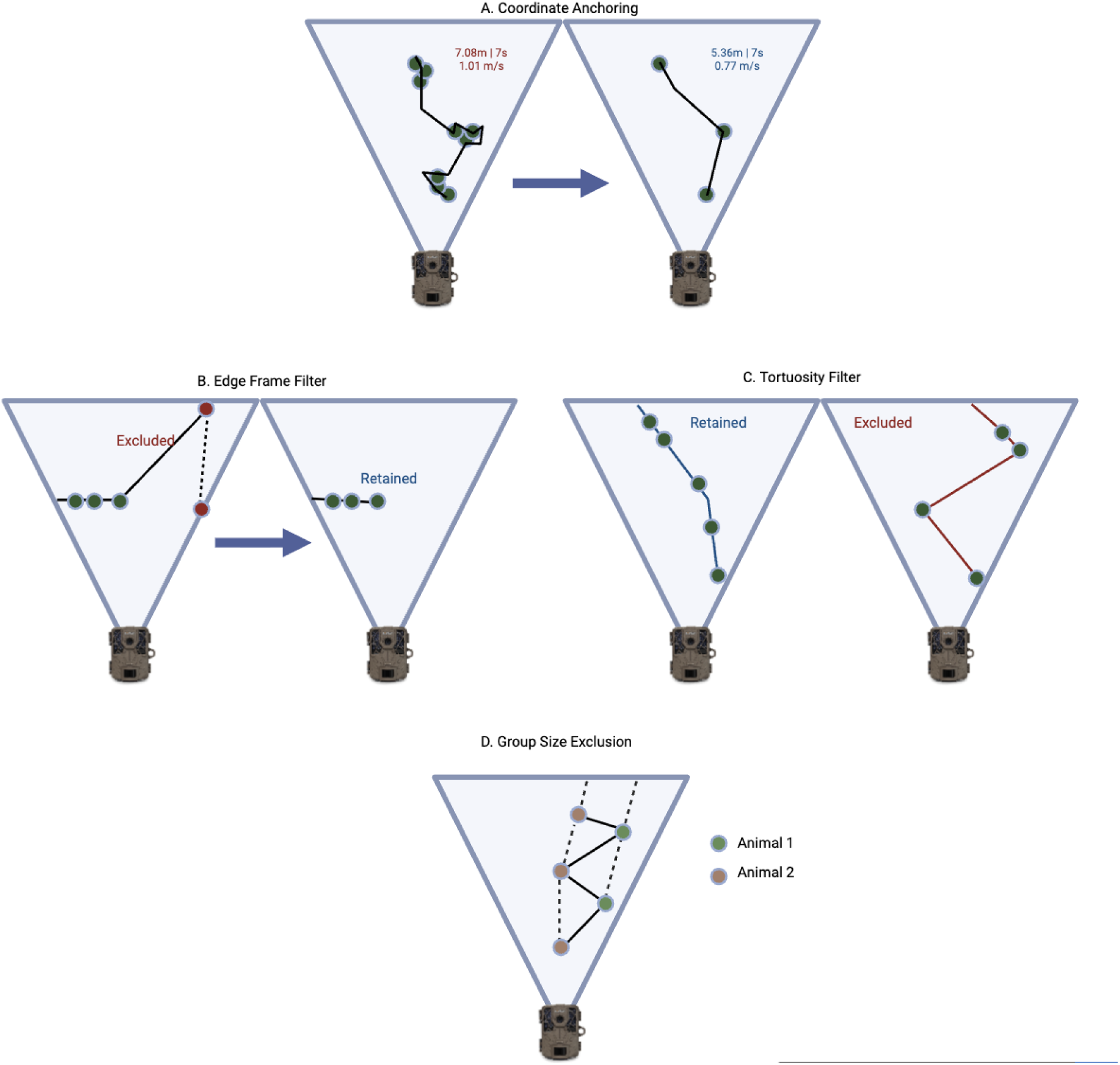
Overview of the filters applied to image-derived animal trajectories prior to speed estimation (A–D). Dashed lines are potential movement paths of animals and solid lines are the AI estimated paths. Frame-level animal coordinates were post-processed to reduce noise and remove unreliable tracks, including anchoring nearby coordinates to limit artificial movement from detection jitter, excluding paths near the edge of the camera viewshed, removing highly tortuous paths likely caused by tracking or distance-estimation error, and excluding sequences with multiple animals where individual identities could not be reliably maintained across frames.

### 2.4 Activity Level Estimation

We calculated activity patterns to convert instantaneous movement speeds into daily movement distances. Activity patterns represent the proportion of each 24-hour period during which animals are active (ω), a critical parameter for estimating day range (Rowcliffe et al. 2014, Palencia et al. 2019).

Activity values were estimated by plotting animal detections across 24-hour periods and applying kernel density smoothing to generate continuous activity curves using the fitact() function in the R package activity (Rowcliffe et al. 2014). The area under the curve represents the proportion of the day animals are active, ranging from 0 (never active) to 1 (continuously active). Under a uniform distribution, an animal would be equally likely to be detected at any hour, corresponding to continuous activity; deviations from this uniform baseline indicate concentrated activity periods, and the area between the fitted kernel density curve and the uniform line quantifies the proportion of the day spent in elevated activity states. This method assumes all individuals in a population follow similar activity patterns, with peak activity times representing periods of maximum movement (Rowcliffe et al. 2014). Uncertainty in activity level estimates was quantified using a bootstrap resampling procedure implemented in the activity package, in which detection times were resampled with replacement across repeated iterations and activity level recalculated each time. The 95% confidence intervals shown reflect the 2.5th and 97.5th percentiles of the resulting bootstrap distribution, with interval width primarily driven by sample size.

Daily movement distance, or day range (DR), was calculated by combining movement speed with activity level as:

*DR (km/day) = Speed (m/s) ✕ Activity (proportion) ✕ 86,400 (seconds/day) / 1,000*

This formula scales instantaneous movement speeds observed during active periods to account for resting and inactive periods throughout the 24-hour cycle.

### 2.5 Staying Time and Group Size

Group size for each sequence was obtained from the Wildlife Insights sequence metadata, which is a manual count of the number of individuals observed per detection event.

Staying time is the total amount of time animals are within the camera’s detection zone. Different density models use staying time differently. For CTDS (Howe et al. 2017) and REST (Nakashima et al. 2018) models they estimate this for each detection regardless of group size (Staying Time groups: ST_g) and then adjust for mean group size elsewhere in the equation. The TIFC method counts per individuals (Staying Time individual: ST_i). Thus two animals in view for 5s would have a ST_g of 5s but a ST_i of 10s. Another nuance is that REST and TIFC literally measure the time animals are in view, while CTDS subsamples the staying time (e.g. ‘snapshot windows’ every 5s). Data for each of these methods can be automatically generated from the bounding boxes generated for animals by the MegaDetector algorithms.

With normal video it would be straightforward to estimate staying time by summing the number of bounding box-seconds. However, typical camera trap image sequences include gaps between triggers, so we created an interpolation method that accounts for gaps in detection within sequences.When cameras operate in burst mode, multiple images can be captured within the same second. To avoid inflating staying time estimates, we first counted the number of bounding boxes per image within each sequence, then adjusted this per unique timestamp. We then constructed a 1-second timeline spanning the full duration of each sequence (first to last timestamp) and populated it with the observed detection count at each second where imagery existed. For seconds within the sequence where no image was captured (gaps between trigger events), we interpolated animal presence using edge detection logic. For short gaps (≤10 seconds) with detections on both ends, we assumed the animal remained present throughout and filled intervening seconds with the representative group size being the mean number of bounding boxes across all detected seconds in the sequence, rounded to the nearest whole number. When the animal’s bounding box appeared at the same frame edge (outer 1/6 of frame width) at both the start and end of a gap, we assumed it had left the detection zone and set intervening seconds to zero. For all other gaps, we linearly interpolated the detection count between the flanking observations and rounded to the nearest whole number, with a minimum of one animal when either endpoint had a detection.

However, if the animal was at the edge of the frame (within the outer 1/6 of the frame width) we recognize they could have walked out of view and then back into view from the same edge. For short gaps (≤10 seconds) with edge detections on both ends, we assumed the animal remained present throughout. For larger gaps (11–59 sec) we assumed it had left the detection zone and re-entered, and thus setting intervening seconds to zero detections. Staying time for each sequence was then calculated as the total number of presence-seconds — the count of seconds during which at least one animal was present (bounding box count > 0). This measure represents the group passage time (ST_g) rather than total animal-seconds: three deer present for 10 seconds yields 10 presence-seconds, not 30. This definition aligns with the REST model formulation (Nakashima et al. 2018) where staying time represents the duration of a single encounter event regardless of group size. For single-frame detections, staying time was set to 1 second (the minimum resolvable interval given the camera’s frame rate).

**Figure 5.**
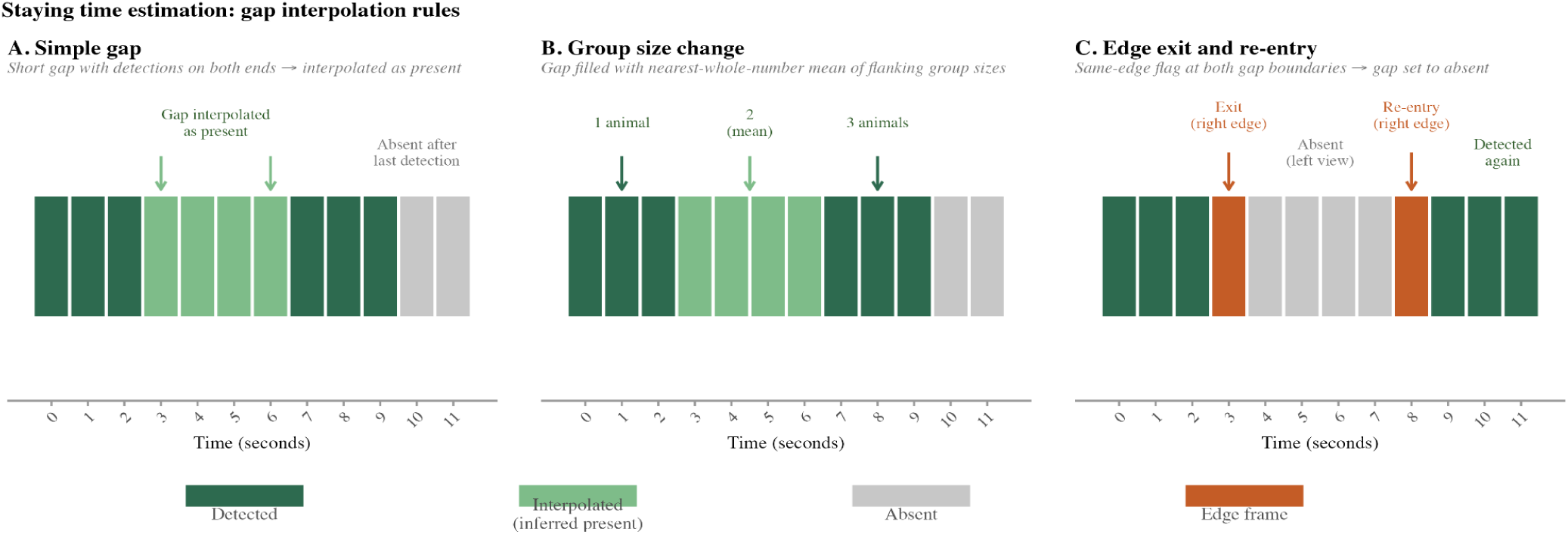
Each second in a detection sequence is classified as detected, interpolated (inferred present), absent, or edge frame. (A) Simple gap: short gaps with detections on both ends are interpolated as present, with absence assigned after the final detection. (B) Group size change: gaps are filled using the nearest whole-number mean of flanking group sizes (e.g., 1 animal → 2 animals → 3 animals). (C) Edge exit and re-entry: when both gap boundaries are flagged as the same edge (e.g., right edge exit and re-entry), the gap is treated as absent, indicating the animal left the field of view before being detected again.

### 2.6 Validation Methods

#### 2.6.1 Distance Estimation Accuracy

We estimated the accuracy and precision of the AI position estimation in two ways: humans and stationary animals. First, we conducted a controlled test where one person stood at a known distance from the camera while another person triggered the camera. We conducted these tests in an open field and in a dense forest to evaluate the effect of vegetation on model performance. The AI pipeline estimated the distance to the person using the same DPT depth model and calibration procedure applied to wildlife images, and we compared AI-estimated distances to true (measured) distances across all test positions. Second, to estimate the precision of the method with actual animal data we looked at variation in the estimated position for instances when an animal was stationary for multiple consecutive images. This allowed us to test the method in day and night images, and to measure errors introduced by changes in animal posture, but not position, which cause small changes in the dimensions of the bounding box.

#### 2.6.2 GPS Telemetry Comparison

Animal tracking data provides movement speed information that can validate our population-level measurements from camera traps. However, direct comparison between camera trap and GPS speeds is complicated by fundamental differences in sampling frequency and path tortuosity. GPS locations collected at typical intervals (30 minutes to several hours) yield straight-line distances that underestimate actual movement, while camera traps sampling at 1.4 fps capture very high resolution movement paths (Rowcliffe et al. 2012, 2016).

To estimate distance moved for GPS tracked animals we applied the ctmm package in R (Calabrese et al. 2016) to publicly available GPS datasets for four species that were also in our camera trap dataset, although from different locations: white-tailed deer, mule deer, coyote, and American black bear. We screened variograms to select only individuals exhibiting home-ranging behavior and fit movement models using the ctmm.select() function, retaining only animals where Ornstein-Uhlenbeck (OU) class models which include both home range confinement and defined velocity processes provided the best fit. Daily movement distances and confidence intervals were calculated using the speed() function, which produces scale-insensitive speed estimates that account for path tortuosity and amount of time between GPS fixes (Noonan et al. 2019).

### 2.7 Effective Detection Distance Estimation

To estimate the effective detection distance (EDD) at each camera deployment, we developed a joint multi-species hierarchical model that borrows strength across species within deployments. This approach improves upon conventional single-species distance sampling by recognizing that all species detected at a given camera share a common detection environment determined by vegetation density, terrain, and camera hardware while differing in their intrinsic detectability due to body size, coloration, and behavior.

#### Detection Data

Distance sampling requires one independent detection distance per encounter event. To avoid inflating the detection histogram with correlated observations from burst photography, we used only the trigger picture — the first frame of each sequence — as the detection distance for each encounter. This ensures each event contributes a single distance observation, consistent with the assumption of independent detections in the distance sampling framework. This approach differs from the formal CTDS protocol described by Howe et al. (2017), which records distances at predetermined ‘snapshot moments’ (typically every 0.25 to 3 seconds) throughout each detection event, generating multiple distance observations per animal pass for as long as the animal remains in the field of view (Howe et al. 2017). Our single-frame approach yields one distance observation per encounter, reducing sample size but ensuring independence among observations. Using all positions at each camera trigger would also be appropriate; standardizing to one observation every 5 seconds for Reconyx and every 3 seconds for Browning could harmonize snapshot intervals across camera models, though we did not implement this here. Detection distances were estimated from the monocular depth output of the Dense Prediction Transformer applied to each trigger image.

We constructed a detection histogram for each species-deployment combination with at least one detection, binning trigger distances into radial distance bins of 3 m width from 0 to B meters, where B was set at 27 m based on inspection of the pooled detection histogram, which showed a sharp decline in detections beyond 26 m consistent with the limits of our calibrated depth range. Each histogram entry L_mj represents the count of detections in distance bin j for observation m, where m indexes a unique species-by-deployment combination.

#### Model Structure

The core of the model decomposes the detection function scale parameter into a deployment-level effect and a species-level offset:

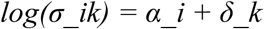

where α_i is a random effect capturing the detection environment at deployment i and δ_k is a species-specific offset reflecting differences in detectability. This decomposition means that data-rich species (e.g., white-tailed deer with 2,245 detections across 138 deployments) inform the site-level detection environment α_i, which in turn improves EDD estimates for data-sparse species at the same camera through partial pooling.

To ensure identifiability, we imposed a sum-to-zero constraint on the species offsets: we estimated K − 1 free offset parameters and derived the Kth as the negative sum of the others. This makes α_i interpretable as the log detection scale for an average species at camera i, and δ_k as each species’ deviation from that average.

#### Covariates on the Deployment Effect

The deployment-level effects were modeled as draws from a normal distribution whose mean is a function of two covariates:

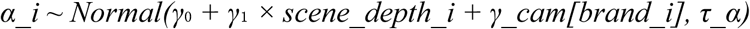

The first covariate, DPT scene depth, is the mean depth value estimated by the Dense Prediction Transformer across the full scene at each deployment, extracted independently of any animal detections. This metric serves as a proxy for openness: deployments in open land cover types (e.g., pasture, cropland, forest with sparse understory) produce larger scene depth values, while those in dense vegetation produce shorter ones. We used the log-transformed, standardized scene depth as the covariate. This approach avoids the circularity of using mean detection distance as a covariate on detection range, because scene depth is measured from the habitat itself rather than from animal detections.

The second covariate, camera brand, captures potential differences in detection capability arising from hardware variation in passive infrared sensor sensitivity, trigger speed, and focal length. Our study used two camera brands (Browning Recon Force Elite and Reconyx Hyperfire), which were randomly assigned to deployments. Camera brand entered the model as a categorical variable with reference-level coding (Browning as reference, γ_cam[1] = 0; Reconyx offset freely estimated).

#### Detection Function

Detection probability at distance d_j for observation m follows a half-normal model modified by a logistic shoulder function:

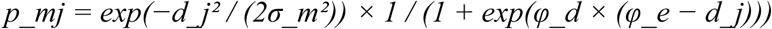

where φ_d and φ_e are shape parameters controlling the near-field detection shoulder. Bin-level detection probabilities are weighted by the relative area of each bin:

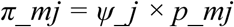

where ψ_j is the fraction of the total survey area (a circle of radius B) covered by bin j. These are normalized to proper probabilities for the multinomial likelihood:

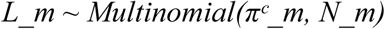

where πᶜ_m is the vector of conditional bin probabilities (normalized to sum to one) and N_m is the total number of detections for observation m. A hierarchical Poisson-Binomial structure models the total detection count at each observation, with deployment- and species-level random effects on expected encounter rates as nuisance parameters.

#### Effective Detection Distance

The EDD for each species-deployment combination was computed from the fitted detection function using a numerical approximation:

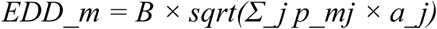

where a_j is the relative area of distance increment j. The EDD represents the radius of a hypothetical perfect-detection circle that would yield the same expected number of detections as the true declining detection function. This is the primary output of the model and serves as input to downstream density estimation.

**Figure 6.**
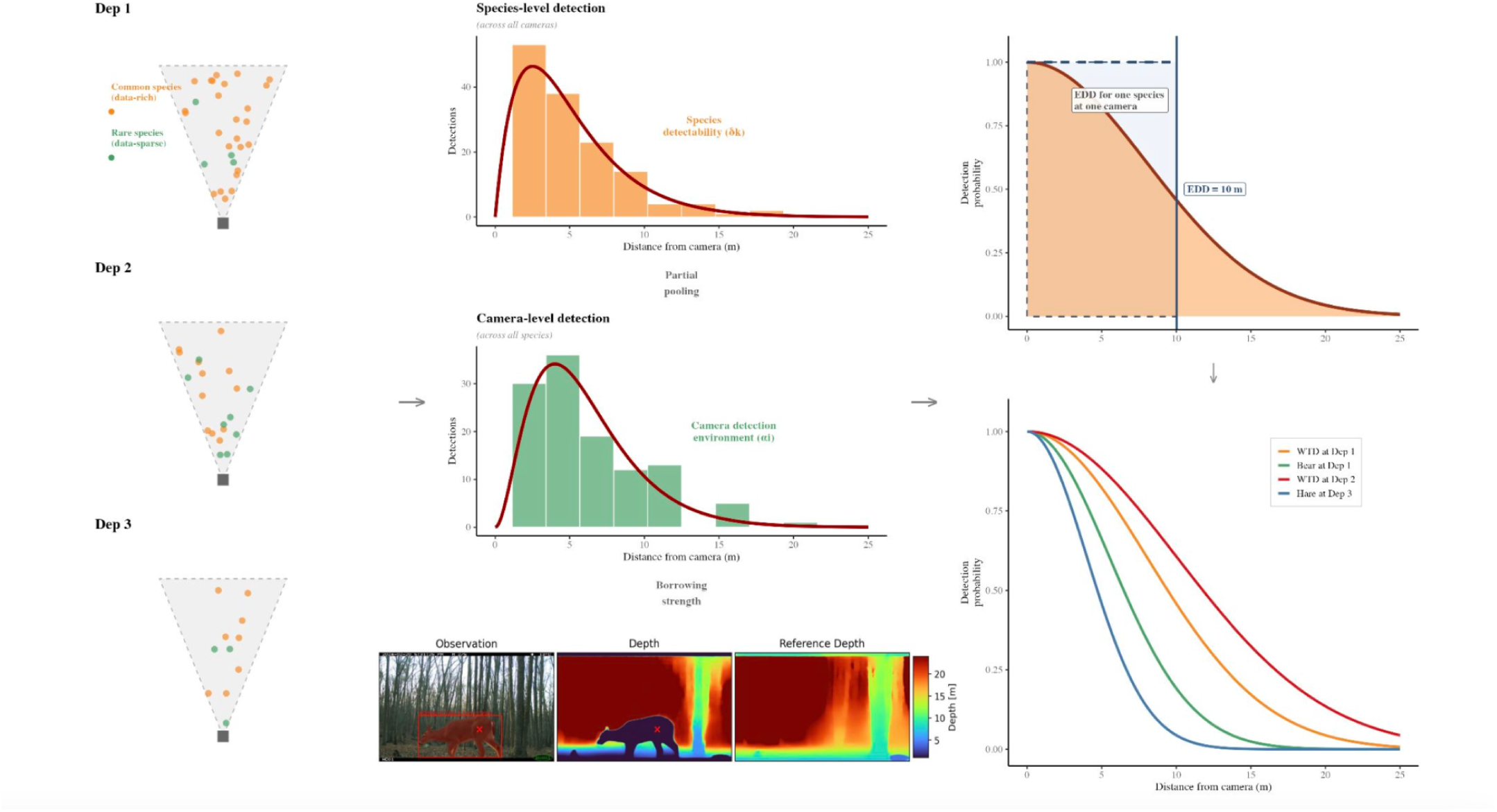
Conceptual diagram of the joint multi-species EDD estimation model. The detection function scale parameter is decomposed into a deployment effect (α_i, green) capturing site-level detection conditions and a species offset (δ_k, orange) capturing differences in detectability. Data-rich species at a camera inform the site effect, improving EDD estimates for rare species at the same camera through partial pooling. Scene depth enters as a covariate on the deployment effect. The final EDD for each species at each camera (right) is derived from the combined detection function.

#### Priors and MCMC

We specified diffuse normal priors (mean = 0, precision = 0.01, equivalent to SD = 10) for all regression coefficients (γ₀, γ₁, γ_cam) and gamma priors (shape = 0.01, rate = 0.01) for precision parameters (τ_α, τ_δ). The logistic shoulder parameters received uniform priors on (0, 10). Species offset parameters were assigned normal priors with precision τ_δ, which was itself estimated, allowing the model to learn the degree of among-species variation in detectability.

We implemented the model in NIMBLE (de Valpine et al. 2017) and ran two MCMC chains of 20,000 iterations each, discarding the first 5,000 as burn-in and thinning by a factor of 10, yielding 3,000 posterior samples for inference. A short diagnostic run (2,000 iterations, 2 chains) was conducted first to verify model initialization and assess preliminary convergence before the full production run. Convergence was assessed using the Gelman-Rubin diagnostic (Brooks and Gelman 1998) and effective sample sizes from MCMCvis (Youngflesh 2018).

#### Covariate Diagnostics

To evaluate whether scene depth was informative for estimating deployment-level detection range, we compared the posterior distribution of γ₁ to its diffuse prior. We quantified informativeness using the precision gain (ratio of prior SD to posterior SD) and assessed whether the 95% posterior credible interval excluded zero. Similarly, we evaluated the camera brand effect (γ_cam[2]) to determine whether hardware differences influenced detection range after accounting for habitat openness.

**Figure 7.**
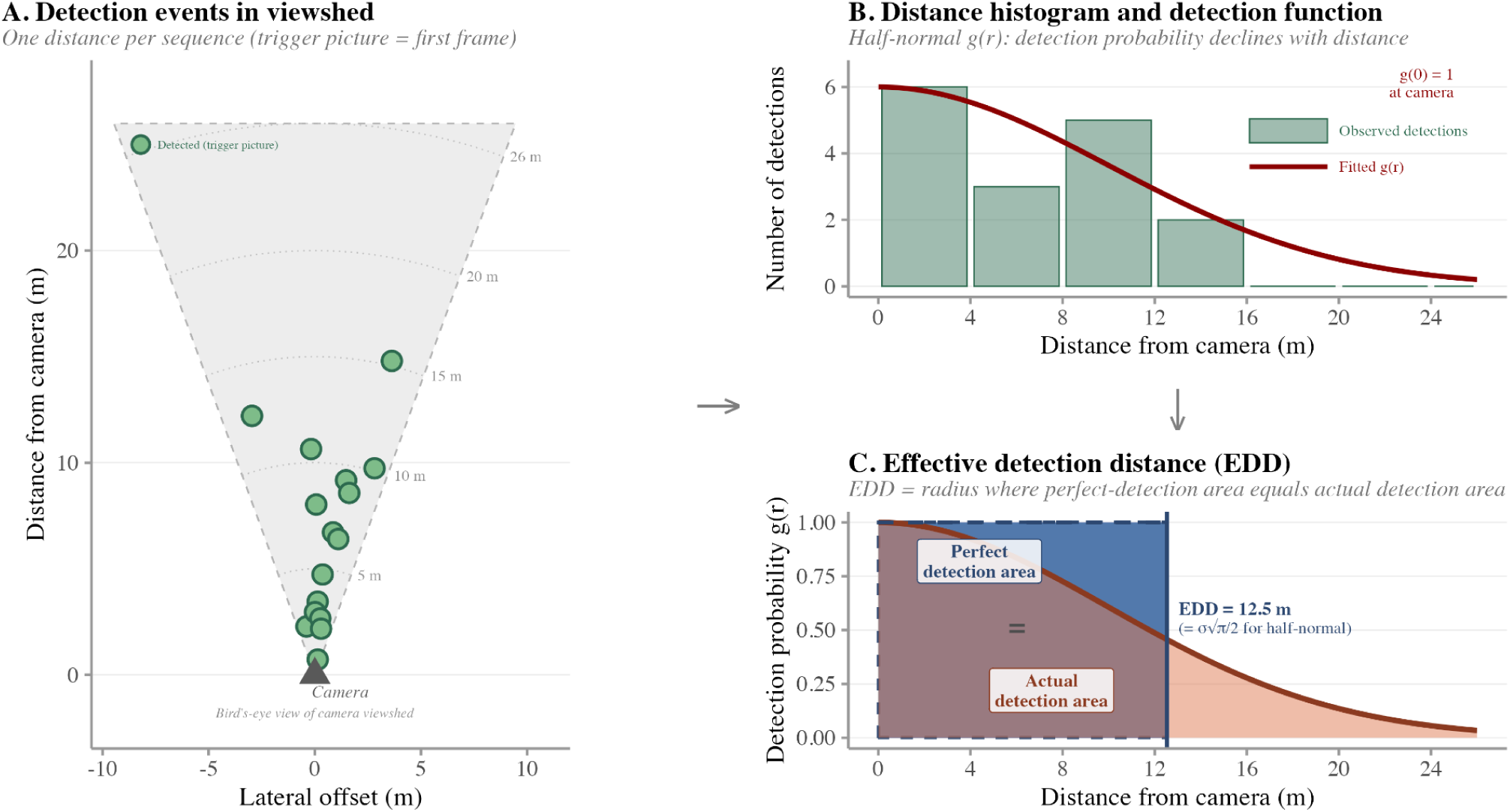
Step-by-step illustration of distance sampling and effective detection distance (EDD) estimation for a single species at a single deployment. (A) Detection events in the camera viewshed. Each point represents one trigger-picture detection plotted at its AI-estimated position (lateral offset and distance from camera). Dashed lines show the camera’s field of view; distance rings mark 5 m intervals. Only the first frame of each sequence (trigger picture) is used to avoid pseudoreplication. (B) Distance histogram and fitted detection function. Trigger-picture distances are binned into 4 m intervals. The fitted half-normal detection function g(r) (red curve) describes how detection probability declines with distance, with g(0) = 1 at the camera. (C) Effective detection distance (EDD). The EDD is the radius at which a circle of perfect detection (blue shaded area, g(r) = 1 out to EDD) would yield the same expected number of detections as the actual declining detection function (red shaded area under the curve). For this example, EDD = 12.5 m, equivalent to σ√(π/2) for the half-normal model.

### 2.8 Density Estimation

To evaluate whether the automated parameter set produces end-to-end density estimates, we applied two density models to white-tailed deer, the most data-rich species in our dataset. The Random Encounter Model (REM; Rowcliffe et al. 2008, 2013) estimates density as:

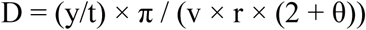

where y/t is the detection rate (detections per unit time), v is day range (km/day), r is effective detection distance (km), and θ is the camera field of view (radians). All four inputs were drawn entirely from the automated pipeline: SBD-corrected movement speed converted to day range via the activity level estimate (day range = speed × activity × 86,400 / 1,000), deployment-specific EDD from the joint hierarchical model, and camera field of view from manufacturer specifications.

We also applied the Random Encounter and Staying Time model (REST; Nakashima et al. 2018), which replaces movement speed with staying time as the key behavioral parameter. REST provides a complementary test of the automated pipeline because it does not require movement speed, instead relying on staying time estimated from bounding box interpolation (Section 2.5) and the detection area defined by camera field of view and detection distance. REST was implemented in NIMBLE with 2,000 iterations, 500 burn-in, and 2 chains.

For both models, detection rate was calculated per deployment as the sum of group sizes across all independent detection events divided by deployment duration in days. We estimated density independently at each of the 138 deployments where white-tailed deer were detected, then averaged across deployments. For REM, uncertainty was quantified using 1,000 bootstrap replicates resampling detection events with replacement, with 90% confidence intervals from the 5th and 95th percentiles. For REST, uncertainty was derived from the NIMBLE posterior distribution.

## III. Results

The SignD model for automated calibration sign detection achieved a recall of 98.7% with perfect precision (1.0) and a mean Average Precision (mAP50) of 99.5% at a 0.50 IoU threshold (mAP50-95 = 81.7%), confirming reliable detection of calibration references across deployments.

In controlled sign validation trials, AI-estimated distances showed a mean absolute error of 2.3 m (95% CI: 1.4-3.1) across 28 test positions spanning 2-29 m from the camera (Figure 8). Accuracy was strongly distance-dependent: MAE was 0.4 m at 0-10 m, 2.1 m at 10-20 m, and 4.4 m beyond 20 m, where AI-estimated depths compressed toward the model maximum. This pattern is consistent with known limitations of monocular depth models at greater range and supports the 26 m maximum distance cutoff applied in subsequent analyses.

**Figure 8.**
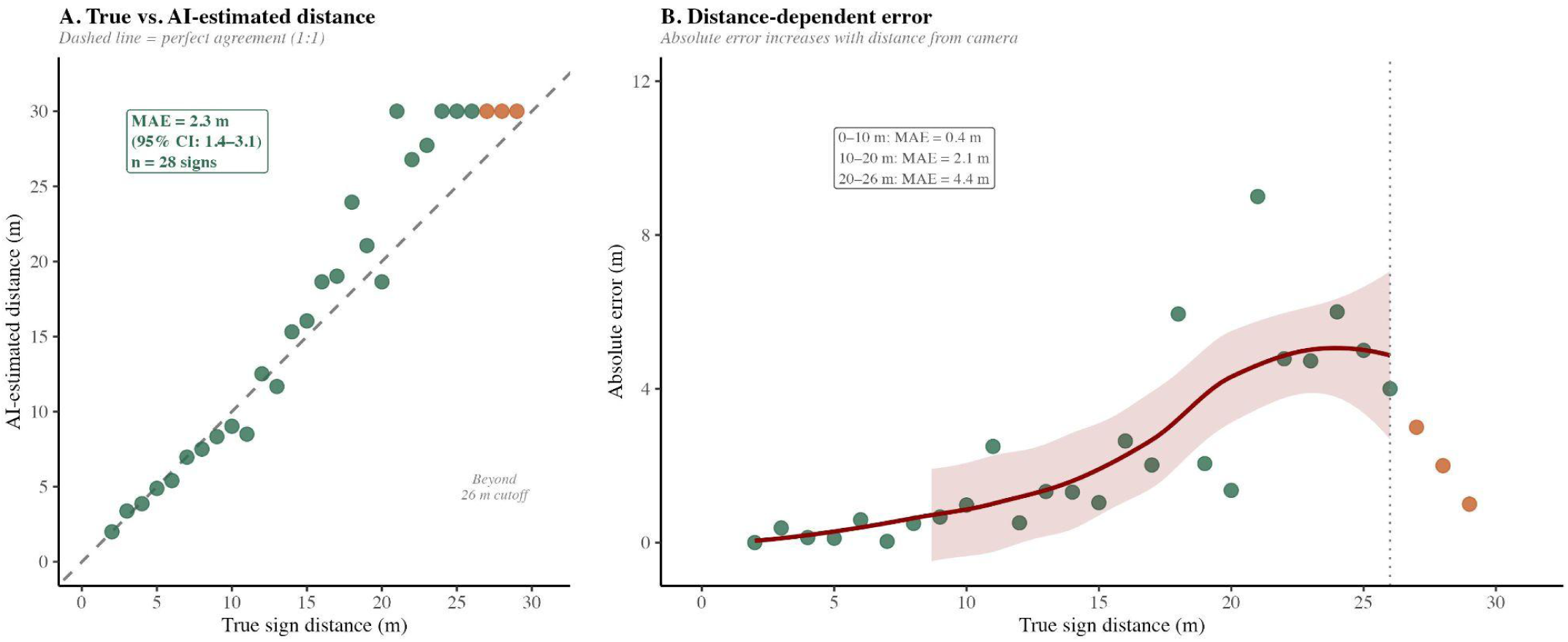
AI distance estimation accuracy validated against calibration signs at known distances (2-29 m). (A) AI-estimated vs. true distance; dashed line = perfect agreement. Estimates compress toward the model maximum beyond ∼20 m. Overall MAE = 2.3 m (95% CI: 1.4-3.1). (B) Absolute error increases with distance: MAE = 0.4 m at 0-10 m, 2.1 m at 10-20 m, 4.4 m at 20-26 m. Green points within the 26 m reliable range; orange beyond. Dotted line marks the 26 m cutoff applied in subsequent analyses.

For stationary animals, within-sequence coordinate jitter (SD of world_z) increased from a median of 0.43 m at 0-3 m to 1.73 m at 15-18 m (Spearman ρ = 0.267, p < 0.001; Appendix S1: Figure S1), reflecting precision limits from posture variation and the progressive loss of informative texture cues at greater range. This distance-dependent noise motivated the species-specific anchoring thresholds applied in speed estimation (Section 2.3).

### 3.2 Movement Speed Estimates

Camera trap analysis yielded SBD speed estimates for 6 species with sufficient data (≥10 sequences; Table 2). Quality filtering required ≥2 frames per sequence and biologically plausible speeds (<20 m/s), retaining sequences with sample sizes ranging from 46 (elk) to 1,021 (white-tailed deer).

**Table 2.** Movement speed estimates derived from camera trap data using the Size-Biased Distribution (SBD) framework, with species-specific anchoring thresholds. n = number of movement sequences retained after quality filtering; Point Anchor setting = species-specific coordinate anchoring threshold below which frame-to-frame displacements are treated as stationary noise (see Appendix S1); SBD Speed = harmonic mean speed corrected for size-biased sampling (Rowcliffe et al. 2016); SE = bootstrap standard error from 1,000 replicates; 95% CI = bootstrap confidence interval (2.5th and 97.5th percentiles). Species with fewer than 10 sequences are excluded.

| Species | n | Point<br>Anchor<br>setting (m) | SBD Speed (m/s) | SE | 95% CI |
| --- | --- | --- | --- | --- | --- |
| White-tailed Deer | 1021 | 0.75 | 0.150 | 0.010 | 0.132–0.170 |
| Mule Deer | 119 | 0.50 | 0.194 | 0.066 | 0.118–0.362 |
| Red Squirrel | 93 | 0.10 | 0.106 | 0.050 | 0.059–0.261 |
| Snowshoe Hare | 90 | 0.10 | 0.109 | 0.024 | 0.079–0.170 |
| American Black Bear | 81 | 0.75 | 0.230 | 0.041 | 0.172–0.329 |
| Elk | 46 | 1.00 | 0.263 | 0.113 | 0.158–0.614 |

### 3.3 Activity Patterns and Day Ranges

Activity patterns were successfully calculated for six species with adequate sampling coverage (n > 40 detections). Activity proportions ranged from 31.2% (snowshoe hare) to 63.9% (American black bear). Ungulates showed moderate to high activity levels: elk (55.0%), white-tailed deer (48.4%), and mule deer (44.7%). Small mammals exhibited lower activity levels, with red squirrels at 35.5% and snowshoe hare at 31.2%.

Activity patterns showed evidence of biological variation among arrays beyond what would be expected from sampling noise alone, with array-level estimates for well-sampled species differing substantially even at sample sizes above 40 detections per site. For example, white-tailed deer activity ranged from 38.7% at arrays with crepuscular patterns to 75.7% at an array exhibiting near-cathemeral activity. Mule deer and red squirrels showed similar cross-array variation (Appendix S1: Figure S1).

Day ranges, calculated by combining SBD speeds with activity patterns, ranged from 2.93 km/day (snowshoe hare) to 12.70 km/day (American black bear) (Table 3). Camera-derived day ranges exceeded GPS ctmm daily distances for all three species with available telemetry data: white-tailed deer (6.26 vs 2.13 km/day, 2.9×), American black bear (12.70 vs 3.28 km/day, 3.9×), and mule deer (7.50 vs 1.74 km/day, 4.3×).

**Table 3.**
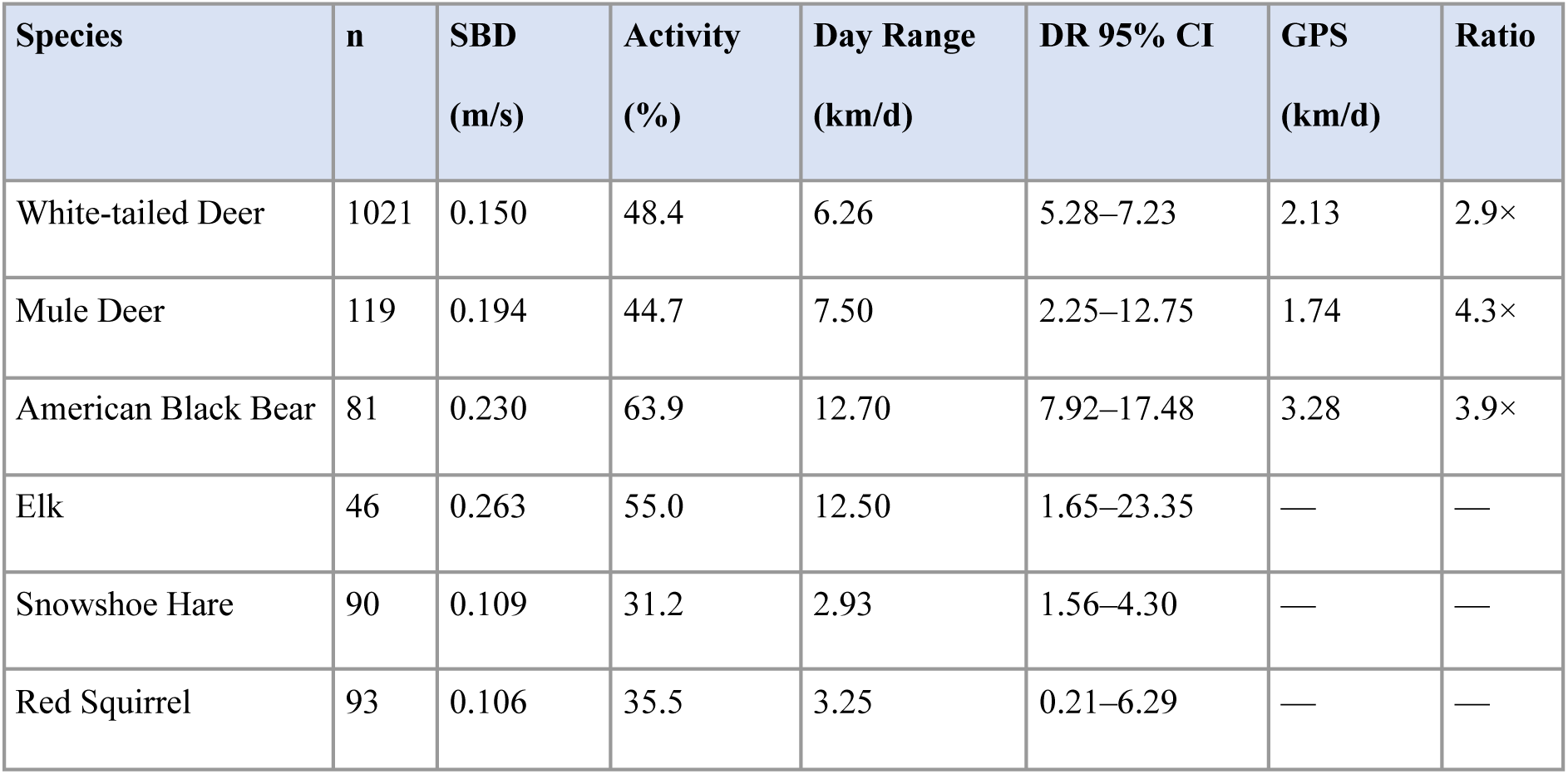
Combined movement parameters: SBD speed, activity level, and day range for species with ≥10 speed sequences. GPS daily distance from ctmm analysis of publicly available telemetry data (Noonan et al. 2019). Camera/GPS ratio shows the factor by which camera-derived day range exceeds GPS-derived daily distance.

| Species | n | SBD<br>(m/s) | Activity<br>(%) | Day Range<br>(km/d) | DR 95% CI | GPS<br>(km/d) | Ratio |
| --- | --- | --- | --- | --- | --- | --- | --- |
| White-tailed Deer | 1021 | 0.150 | 48.4 | 6.26 | 5.28–7.23 | 2.13 | 2.9× |
| Mule Deer | 119 | 0.194 | 44.7 | 7.50 | 2.25–12.75 | 1.74 | 4.3× |
| American Black Bear | 81 | 0.230 | 63.9 | 12.70 | 7.92–17.48 | 3.28 | 3.9× |
| Elk | 46 | 0.263 | 55.0 | 12.50 | 1.65–23.35 | — | — |
| Snowshoe Hare | 90 | 0.109 | 31.2 | 2.93 | 1.56–4.30 | — | — |
| Red Squirrel | 93 | 0.106 | 35.5 | 3.25 | 0.21–6.29 | — | — |

**Figure 9.**
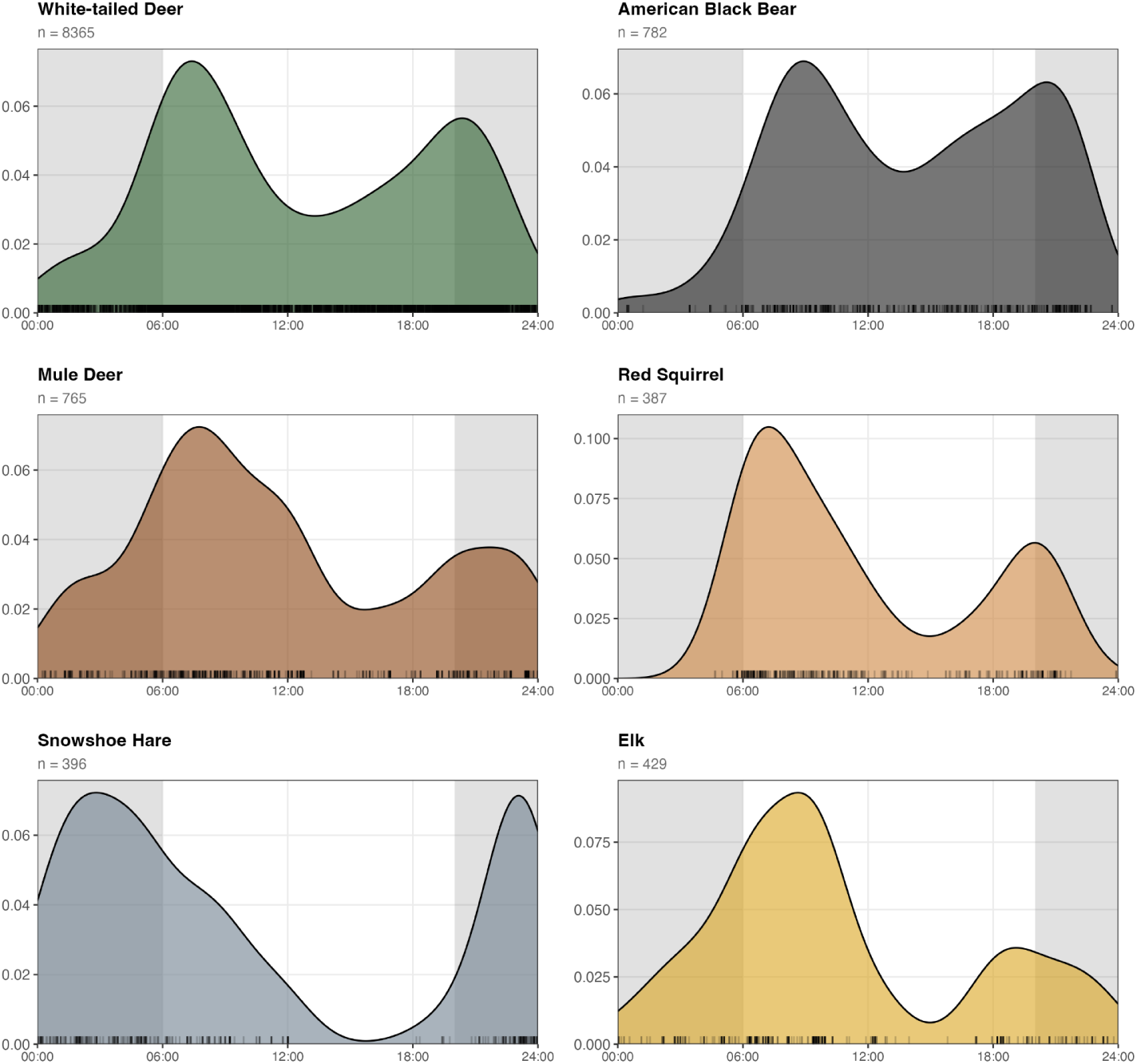
Diel activity patterns for six species. Kernel density activity curves plotted over the 24-hour cycle (0:00–24:00) for each species with sufficient detections (n ≥ 15). Tick marks along the x-axis show individual detection times. The shaded area under each curve represents the estimated proportion of the day spent active (activity level) with 95% confidence intervals. Species are arranged to highlight contrasting diel strategies: crepuscular (white-tailed deer, mule deer), diurnal (red squirrel), nocturnal (snowshoe hare), and broadly active (American black bear).

### 3.4 Staying Time

Mean staying times (ST_g) ranged from 3.8 seconds (red squirrel) to 36.8 seconds (elk) after correction for burst photography artifacts (Table 4). Distributions were strongly right-skewed for all species, with medians substantially lower than means, reflecting the mix of brief transits and prolonged stationary events (bedding, foraging in place) captured by cameras.

**Table 4.**
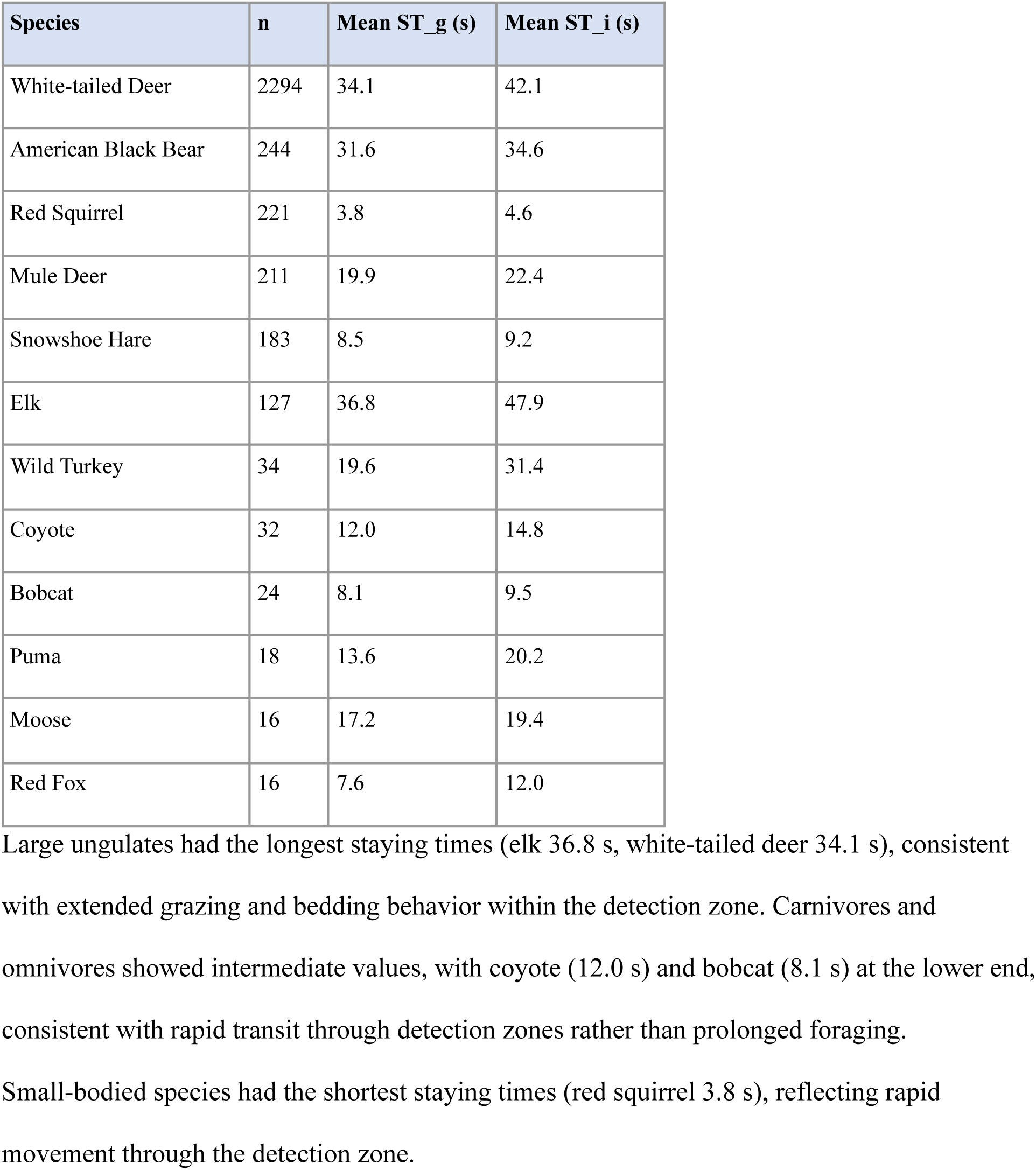
Staying time estimates by species. ST_g = group presence-seconds (time any animal present); ST_i = individual animal-seconds (summing across all individuals per second).

| Species | n | Mean ST_g (s) | Mean ST_i (s) |
| --- | --- | --- | --- |
| White-tailed Deer | 2294 | 34.1 | 42.1 |
| American Black Bear | 244 | 31.6 | 34.6 |
| Red Squirrel | 221 | 3.8 | 4.6 |
| Mule Deer | 211 | 19.9 | 22.4 |
| Snowshoe Hare | 183 | 8.5 | 9.2 |
| Elk | 127 | 36.8 | 47.9 |
| Wild Turkey | 34 | 19.6 | 31.4 |
| Coyote | 32 | 12.0 | 14.8 |
| Bobcat | 24 | 8.1 | 9.5 |
| Puma | 18 | 13.6 | 20.2 |
| Moose | 16 | 17.2 | 19.4 |
| Red Fox | 16 | 7.6 | 12.0 |

### 3.5 GPS Telemetry Comparison

Continuous-time movement model (ctmm) analysis of GPS tracking data from publicly available datasets provided daily movement distance estimates for four focal species. Camera-derived day ranges exceeded ctmm-derived GPS daily distances consistently: white-tailed deer (2.9×), American black bear (3.9×), and mule deer (4.3×). These ratios are presented alongside camera-derived day ranges in Table 4 above.

### 3.6 Effective Detection Distance Estimation

The joint multi-species model estimated EDD for 480 species-by-deployment combinations across 9 species and 181 deployments, using 3,284 trigger pictures. Mean EDD across species ranged from 4.07 m (red squirrel) to 7.08 m (elk), with substantial within-species variation across deployments reflecting differences in vegetation density, camera placement, and local detection conditions (Table 5; Figure 10).

**Figure 10.**
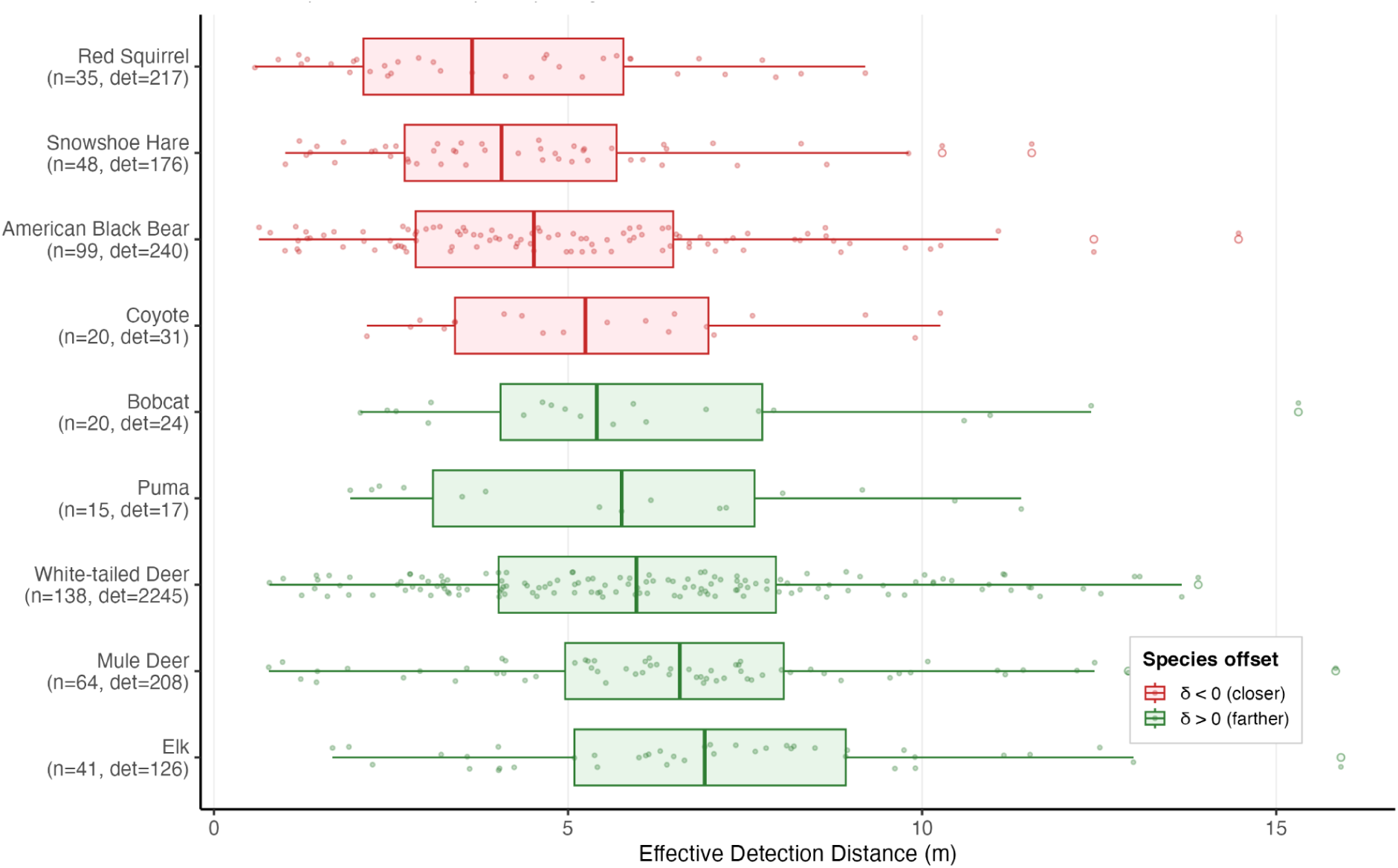
Deployment-level effective detection distance (EDD) estimates by species from the joint multi-species hierarchical model with partial pooling. Each point represents the posterior mean EDD for one species at one deployment. Boxplots summarize the distribution across deployments. Species are colored by their estimated species offset (δ_k): red indicates species detected at shorter distances than the mean across all species at that deployment (δ < 0) and green indicates species detected at greater distances (δ > 0). Species are ordered by median EDD. Sample sizes indicate the number of deployments with detections (n) and total trigger-picture detections (det) for each species.

Species offsets (δ_k) from the joint model revealed that larger-bodied ungulates were detected at greater distances than average, while small-bodied species and ground-dwelling birds had shorter detection ranges. White-tailed deer (δ = +0.095, 95% CI: 0.044 to 0.146), elk (δ = +0.093, CI: 0.008 to 0.185), and mule deer (δ = +0.088, CI: 0.009 to 0.172) had the largest positive offsets, consistent with their larger body sizes increasing detection probability at greater distances. The overall standard deviation of species offsets (SD = 0.097) was substantially smaller than the deployment-level random effect (SD = 0.297), indicating that site-to-site variation in detection conditions exceeded species-level differences in detectability by approximately 3:1. This confirms that deployment-level habitat effects dominate the detection process, supporting the joint model’s approach of sharing deployment information across species.

#### Covariate Effects

The scene depth covariate (γ₁), our CV derived measure of viewshed, had a strongly positive effect on detection range: posterior mean = 0.266 (95% CI: 0.215 to 0.317), with the credible interval clearly excluding zero. The precision gain from prior to posterior was 390-fold (prior SD = 10, posterior SD = 0.026), indicating that the data were highly informative for this parameter. Deployments with greater scene depth (more open habitat) had correspondingly larger detection scales, consistent with increased detectability in less cluttered environments.

The camera brand effect was not statistically distinguishable from zero. The Reconyx offset relative to Browning (reference) was −0.060 (95% CI: −0.164 to 0.042), indicating no significant difference in detection range between the two camera brands after accounting for habitat openness. On the exponential scale, this corresponds to Reconyx cameras having approximately 94% of the detection scale of Browning cameras, a difference that is both small and uncertain.

Model convergence was confirmed by Gelman-Rubin diagnostics near 1.0 for key parameters (γ₁ R-hat = 1.00, sd_alpha R-hat = 1.00) and adequate effective sample sizes (n.eff = 720 for γ₁, 362 for sd_alpha, 567 for sd_delta).

### 3.7 Density Estimation

To demonstrate end-to-end functionality, we computed density estimates for white-tailed deer using two models with fully automated parameter inputs. The REST model estimated white-tailed deer density at 11.6 animals/km² (90% CI: 9.7–13.8). The REM, which requires movement speed as an input, estimated 21.4 animals/km² (90% CI: 20.9–22.0). Note that REM CI reflects bootstrap variability over deployments only, not parameter uncertainty in EDD/speed/activity. Both estimates are within an order of magnitude of published white-tailed deer densities for forested habitat in the inland Northwest (Wiegers et al. 2025), though the REM estimate is higher than most published values for comparable montane forest habitat. The difference between the two models, which shared the same automated detection parameters but differed in their movement input, suggests that the speed parameter warrants further investigation, as discussed below.

## IV. Discussion

### Overview: automation unlocks scale, scale unlocks new capabilities

Our AI-driven computer vision pipeline processed 122,574 frames from 181 deployments across Washington and Montana, automating extraction of the full suite of parameters required for camera trap density estimation for the mammal community at our study sites. Complete parameter sets (speed, activity, staying time, and effective detection distance) were generated for six species with sufficient sample sizes, and partial estimates for an additional six species, without manual annotation at any stage. This automation enabled three capabilities that would be impractical with manual methods: deployment-specific effective detection distances across hundreds of cameras, community-wide speed estimation using the size-biased distribution framework, and automated behavioral parameters including activity levels and staying times. The most important finding was that deployment-level variation in effective detection distance dwarfed species-level variation by approximately 3:1, demonstrating that the area surveyed by each camera varies massively across sites and is likely the most consequential detection parameter for density estimation. Speed estimates validated against GPS telemetry confirmed that camera-derived encounter velocities are systematically higher than landscape-scale displacement rates, a difference that reflects sampling scale rather than measurement error but requires careful interpretation. A series of quality control filters proved necessary to make

AI-derived coordinates usable for parameter estimation, retaining approximately 92.5% of frame-level detections and 56.2% of movement sequences after filtering. These results establish a foundation for fully automated, community-level density estimation across large camera trap networks (Rowcliffe et al. 2008, Howe et al. 2017, Wearn et al. 2022), a capability increasingly needed as standardized camera trap programs expand across cities (Magle et al. 2019), countries (Rooney et al. 2025), and continents (Kays et al. In Review).

This work makes four contributions to camera trap density estimation. First, we present the first fully automated pipeline for extracting all four parameters required by camera trap density models (effective detection distance, movement speed, activity level, and staying time) directly from imagery, without manual annotation. Second, the joint multi-species hierarchical detection model provides deployment-specific EDD estimates by decomposing detectability into site-level and species-level components, revealing that site conditions dominate detectability by 3:1 over species identity. Third, validation against GPS telemetry clarifies that camera-derived speeds measure fine-scale encounter velocity rather than landscape-scale displacement, a distinction with direct implications for how speed is used in density models. Fourth, end-to-end density estimation using REST produced plausible white-tailed deer estimates, demonstrating that the automated parameter set can support applied density estimation without manual measurement at any stage.

### Camera survey area

#### Joint EDD model: the key methodological advance

The area surveyed by each camera is probably the largest source of variation in detectability, a fact unsurprising to any experienced camera trapper, but critical to quantify and never previously accomplished at this scale. Our joint multi-species hierarchical model estimated effective detection distances for 480 species-by-deployment combinations across 9 species and 181 deployments, a scope only possible with automated distance estimation. Deployment-level variation in the detection scale parameter (SD = 0.297) exceeded species-level variation (SD = 0.097) by approximately 3:1, meaning that site-to-site differences in vegetation density, terrain, and camera placement dominate the detection process far more than species identity. This finding has a critical implication: the standard practice of applying a single EDD per species across an entire study introduces substantial uncertainty whenever cameras span heterogeneous environments. The joint model addresses this by decomposing the detection function into shared deployment effects and species-specific offsets, allowing data-rich species such as white-tailed deer (2,245 detections across 138 deployments) to inform detection conditions at cameras where rare species have few observations. This represents a substantial advance over conventional distance sampling approaches (Buckland et al. 2001, Howe et al. 2017, Rowcliffe et al. 2011) and offers a mechanistically interpretable detection parameter that is directly transferable to density estimation.

Species offsets from the joint model confirmed expected body-size effects on detectability. Larger-bodied ungulates (white-tailed deer δ = +0.095, elk δ = +0.093, mule deer δ = +0.088) were consistently detected at greater distances, while small-bodied species (red squirrel, snowshoe hare) and American black bear had negative offsets. The overall standard deviation of species offsets (SD = 0.097) was modest relative to deployment-level variation, reinforcing that habitat structure rather than species identity drives most variation in detection range. However, species offsets were precisely estimated and biologically interpretable, suggesting that the joint model successfully separates the two sources of variation.

Any distance sampling approach is an improvement over occupancy-based detection probability applied to camera traps (MacKenzie et al. 2002), which simply uses re-detection across time periods at the same site as an indication of detectability. For camera traps this variation reflects primarily whether an animal happened to pass within camera range during a given interval, and thus is more of a measure of availability rather than detectability in any mechanistic sense.

#### Scene depth drives EDD; hardware does not

The scene depth covariate, an AI-derived measure of openness extracted from the same depth estimation pipeline, strongly predicted deployment-level detection range. The precision gain from prior to posterior was 390-fold, confirming that the data were highly informative for this parameter. Deployments with greater scene depth, corresponding to more open areas, had correspondingly larger detection scales, consistent with the well-established relationship between vegetation structure and camera detection range (Hofmeester et al. 2017). This validates AI-derived vegetation metrics as ecologically meaningful predictors of detection range and provides a non-circular approach to modeling detectability, since scene depth is measured from the environmental conditions specific to the camera location rather than from animal detections. The camera brand effect was not statistically distinguishable from zero. However, this negligible hardware effect may partly reflect our study’s dense forest environments, where vegetation rather than sensor range limits detection distance. In more open areas where maximum detection distance is set by sensor capability rather than vegetation, camera brand differences could become more consequential in EDD (Trolliet et al. 2014, Meek and Pittet 2012), and thus should be included in any application of this distance sampling model.

### Animal movement and behavior

#### Speed estimation: promise and heterogeneity

Size-biased distribution speeds were estimated for 6 species with sufficient sample sizes, ranging from 0.109 m/s (snowshoe hare) to 0.263 m/s (elk). Within-species speed estimates varied substantially across sites: the coefficient of variation across arrays was 94.7% for white-tailed deer, 88.6% for mule deer, and 79.7% for red squirrel, likely reflecting site-specific differences in habitat structure, behavioral contexts, and the proportion of foraging versus transit movements captured at each location. Site-specific speed estimates would be preferable to study-wide means where sample sizes allow; our current deployment durations and camera densities did not consistently yield sufficient sequences per array, but longer deployments and larger arrays would enable array-level estimation. Notably, our pipeline produced the first camera-derived day range estimates for several small-bodied species, including red squirrel (3.25 km/day) and snowshoe hare (2.93 km/day), that have not been tracked with GPS collars, filling a gap in movement ecology for species where telemetry is impractical.

#### Camera vs. GPS speeds measure different things

Camera-derived day ranges exceeded ctmm GPS estimates for all three species with available telemetry data: white-tailed deer (2.9×), American black bear (3.9×), and mule deer (4.3×). The GPS datasets available for comparison were not ideal, coming from different regions and time periods with relatively low fix intervals, which limits direct inference (Morin et al. 2022). However, the systematic direction of the discrepancy is expected. Camera traps sampling at approximately 1.4 fps capture near-complete movement paths including turns, foraging movements, and other fine-scale behaviors, while GPS telemetry at hourly intervals primarily measures net displacement between fixes. This difference in resolution makes higher camera-derived distances unsurprising: when GPS fix intervals substantially exceed the velocity autocorrelation timescale, velocity information decays entirely between fixes and no estimator can recover true path length from position data alone (Gunner et al. in press, Calabrese et al. 2016). Gunner et al. establish that τv ranges from roughly 12 to 54 seconds across species, with most falling in the 20–40 second range; at hourly fix intervals, sampling occurs far beyond 3τv for virtually any terrestrial species, meaning continuous-time movement models default to estimating displacement from position autocorrelation and home range structure rather than actual path velocity. Camera traps operating at approximately 1.4 fps resolve movement well below τv and capture the full step-turn structure of the path through the frame and are not inflated estimates, but reflect fine-scale tortuosity that hourly GPS telemetry cannot see regardless of which estimator is applied.

The speed measure needed for REM density estimation is encounter velocity, the fine-scale instantaneous speed at which animals move through the landscape and cross camera detection zones. Despite this, past REM studies relying on low-resolution GPS have produced plausible density estimates (Rowcliffe et al. 2008, Henrich et al. 2024, Palencia et al. 2021), which likely reflects spatio-temporal covariation in movement speed (Broadley et al. 2019). Alternatively, if GPS underestimates are consistent enough across species and sites then other REM error sources (e.g. detection area) might dominate the uncertainty budget. Camera-derived encounter velocity sidesteps this problem entirely, measuring speed at the scale and behavioral context directly relevant to detection. More high-resolution GPS tracking data, particularly from the same populations being camera-trapped, would help calibrate the camera-GPS speed relationship and clarify how movement speed scales across measurement resolutions.

#### Staying time: behavioral signal and density implications

Movement rate is also reflected in staying time: species with slower movement through the detection zone had correspondingly longer staying times. Mean staying times ranged from 3.8 seconds (red squirrel) to 36.8 seconds (elk), with strongly right-skewed distributions for all species reflecting the mix of brief transits and prolonged stationary events captured by cameras. Large ungulates had the longest staying times, consistent with extended grazing within the detection zone. Full video would provide more precise staying time estimates by eliminating the need for gap interpolation; our method innovates by constructing a continuous 1-second timeline from burst imagery and using edge detection logic to infer presence during gaps. Our bounding box interpolation approach to staying time estimation differs from the snapshot-based survey duration methods used in formal Camera Trap Distance Sampling (CTDS). Kühl et al. (2023) demonstrated that camera recovery time and retrigger delay reduce the effective number of snapshot moments during a survey. They addressed this by estimating mean inter-trigger intervals for each camera model and using these to calculate effective survey duration, which replaces the nominal survey duration in the CTDS effort calculation. Our approach sidesteps this issue entirely: by constructing a continuous 1-second timeline from burst imagery and interpolating gaps using edge detection logic, we directly estimate the duration of each encounter event without relying on predefined snapshot intervals or needing to account for camera-specific recovery characteristics. One limitation is that prolonged stationary behaviors such as bedding or sleeping break our gap-filling assumptions. Future AI models capable of recognizing sleeping or bedded animals could improve staying time accuracy for species with long resting bouts within the detection zone. For species where staying time includes substantial resting, we presume our estimates do not capture all sleep events and therefore adjust by the proportion of the day active.

#### Activity patterns: biological variation across sites

Activity levels were estimated with sufficient precision for six species with ≥40 detections. Array-level variation exceeded sampling noise for well-sampled species: white-tailed deer activity ranged from 38.7% at arrays with classic crepuscular patterns to 75.7% at an array exhibiting near-cathemeral activity. Similar cross-array variation was observed for mule deer and red squirrel (Appendix S1: Figure 2). Study-wide pooling of activity estimates is acceptable for the present parameter-estimation study, but our pipeline enables site-specific estimates when sample sizes are sufficient (≥40 detections per species per array), which would better capture local behavioral patterns for density estimation applications. Seasonal changes in activity patterns also likely occur and represent an additional source of variation warranting further investigation (Frey et al. 2017).

### Limitations and future work

While computer vision approaches greatly increase our sample size overall, we were still unable to estimate parameters for some species. Longer deployments would increase sample sizes but introduce concerns about population closure and seasonal behavioral changes. Deploying more cameras or incorporating data from publicly available data such as Snapshot USA (Rooney et al. 2025) could address sample size limitations while maintaining temporal closure.

Depth uncertainty propagates through multiple stages of the pipeline: coordinate errors affect frame-to-frame displacements, which affect SBD speed estimates, which in turn affect day range and density calculations. We do not formally quantify this propagation, meaning that the confidence intervals reported for speed and density reflect sampling variability but not measurement uncertainty. Future work could address this through simulation-based error propagation, where known perturbations are added to AI coordinates and their effects on downstream parameter estimates are quantified, or through paired validation studies comparing AI-derived coordinates against high-precision reference measurements (e.g., LiDAR or stereo camera systems).

Several technical improvements could reduce the need for filtering in future implementations. Improved depth estimation architectures (e.g., Depth Anything V2; Niccoli et al. 2025), higher frame-rate video enabling reliable individual tracking within groups, and additional calibration points or self-calibrating approaches would improve coordinate precision and reduce reliance on post-hoc filtering. Animals may also move at different speeds in groups versus alone, introducing potential bias in speed estimates from sequences where group size could not be determined.

### Path to fully automated density estimation

All parameters required for Random Encounter Model (Rowcliffe et al. 2008), Camera Trap Distance Sampling (Howe et al. 2017), the Random Encounter and Staying Time model (Nakashima et al. 2018), the Space-to-Event model (Moeller et al. 2018), and the Time-to-Event model (Moeller et al. 2018) are now automatable: effective detection distance, movement speed, activity level, and staying time. The REST and REM density estimates for white-tailed deer (11.6 and 21.4 animals/km², respectively) provide an initial test of the automated pipeline for density estimation. Both estimates are within an order of magnitude of published densities for the region, and the REST estimate in particular appears plausible for productive forested habitat. REM produced a higher estimate, which may reflect the sensitivity of speed-dependent models to variation in movement rates across sites, seasons, measurement scales, and assumptions about the proportion of the day spent active. Camera traps sample movement at sub-second resolution within a small detection zone, while published GPS-derived speeds used to parameterize REM in other studies are typically measured at much coarser temporal grain. How movement speed scales across these measurement resolutions remains an open question. Recent work compiling movement traits across species and populations (Beumer et al. 2026) highlights substantial variation in movement rates across sites and seasons, underscoring the importance of measuring speed from the same populations and time periods being surveyed, which is precisely what the camera-based approach enables. More broadly, variation in speed between sites, seasons, camera technology, measurement scale, and activity assumptions may make REM more sensitive to parameter uncertainty than models like REST that do not require movement speed as an input.

Our density comparison showed that REST, which uses staying time rather than movement speed, produced a more plausible estimate than REM for white-tailed deer, suggesting that the speed parameter introduces more uncertainty than other automated inputs. The three remaining automated parameters — deployment-specific EDD, activity level, and staying time — produced plausible density estimates through REST and are immediately usable as inputs for density models. The joint EDD model in particular addresses what may be the most consequential source of uncertainty in camera trap density estimation: unaccounted site-to-site variation in detection area. As AI continues to transform wildlife monitoring (Tuia et al. 2022), fully automated density estimation across large camera networks is now within reach, a capability essential for meeting global biodiversity monitoring targets and improving the taxonomic and geographic coverage of population trend databases such as the Living Planet Index (Loh et al. 2005) and TetraDENSITY (Santini et al. 2018).

## Code Availability

All R code for the parameter estimation pipeline, including the joint multi-species hierarchical effective detection distance model (NIMBLE), quality filtering, movement and staying-time estimation, and density estimation, is archived on Zenodo (https://doi.org/10.5281/zenodo.20573173) and maintained at https://github.com/sierramcmurry/cameratrap-density-parameter-pipeline. The AI depth estimation pipeline (MegaDetector, Segment Anything Model, and Dense Prediction Transformer) that generates the input coordinates is archived on Zenodo (https://doi.org/10.5281/zenodo.20603310) and maintained at https://github.com/Alyetama/distance-estimation. Processed data supporting the analyses, including per-deployment DPT scene-depth values (deployment_scene_depth.csv), are archived in a separate Zenodo (https://doi.org/10.5281/zenodo.20645128). All repositories are publicly available without embargo. Together they provide the full workflow from raw camera trap imagery to density parameter estimates.

