## Appendix for "Automated Parameter Estimation for Camera Trap Density Models Using Computer Vision-Enhanced Distance Sampling"

Ecological Monographs

### Appendix S1

This appendix provides supporting methods, tables, and figures referenced in the main text.

#### Section S1: Field protocol for camera calibration

Calibration suggestions for field technicians, with example reference images illustrating the calibration setup for a single camera deployment. A high-contrast distance sign was sequentially positioned at known distances along the camera's viewing axis to provide deployment-specific reference imagery.

**Calibration tips for field technicians:**

- Mark out positions at 3-m intervals from the camera with measuring tape or phone app.
- Arm camera for detection before starting.
- Position at 3-m mark to begin.
- Hold sign at waist height.
- Trigger capture via hand wave (two-person team) or lateral movement (solo).
- Wait 5 seconds (team) or 10 seconds (solo) between measurements.
- Place used signs behind unused ones to prevent field-of-view contamination.
- Continue at 3-m intervals until beyond maximum detection range.

#### Section S2: Species-specific coordinate anchoring threshold selection

##### S2.1 Threshold evaluation

We evaluated anchoring thresholds ranging from 0.1 m to 1.5 m and confirmed that a uniform threshold introduced selection bias across body sizes. At a uniform 0.5 m anchoring, red squirrels showed a 642% artificial increase in SBD speed due to exclusion of slow foraging sequences where animals moved short distances between frames.

##### S2.2 Kneedle algorithm results

The Kneedle algorithm (Satopää et al. 2011) produced reliable thresholds for seven species: American black bear (0.75 m), bobcat (1.0 m), elk (1.0 m), mule deer (0.3 m), red squirrel (0.1 m), white-tailed deer (0.3 m), and wild turkey (0.5 m). For four species the algorithm was flagged as unreliable: coyote, moose, and puma had fewer than 10 sequences surviving at the detected elbow and reverted to body-size defaults (0.3 m for coyote and puma, 0.5 m for moose); snowshoe hare showed an inverse curve where speed increased with threshold and reverted to the default (0.1 m).

##### S2.3 Manual overrides

After reviewing the sweep results, we manually overrode two species where the Kneedle suggestion did not capture the full noise floor. For white-tailed deer, the algorithm identified 0.3 m as the elbow, but SBD speed continued decreasing through 0.75 m (0.157 m/s vs. 0.187 m/s at 0.3 m), indicating that postural noise from head turns and weight shifts in this large-bodied species extends beyond 0.3 m; we raised the threshold to 0.75 m. For mule deer, the algorithm suggested 0.3 m, but the sweep showed SBD speed continued decreasing to a minimum at 0.5 m before rising at higher thresholds; the Kneedle detected the elbow prematurely, so we overrode to 0.5 m. We also raised the tortuosity threshold for elk from 3 to 4, as this large-bodied species exhibited genuinely longer path-to-displacement ratios within detection zones (Figure S2).

##### S2.4 Final thresholds

Species-specific coordinate anchoring thresholds and the selection method for each species are reported in Table S5.

#### Tables

**Table S1.** Camera model specifications including focal length, horizontal and vertical field of view, and sensor resolution for each camera model used. Specifications compiled from manufacturer documentation. H-FOV = horizontal field of view; V-FOV = vertical field of view.

| Camera model | Focal length (mm) | H-FOV (°) | V-FOV (°) | Sensor (MP) |
|---|---|---|---|---|
| Browning Recon Force Elite | 1.82 | 41.0 | 30.0 | 20 |
| Reconyx Hyperfire 2 | Not published | 38.2 | 29.1 | 3 |
| Reconyx HC500 Hyperfire | Not published | 42.2 | 32.3 | 3 |

**Table S2.** Site-level size-biased distribution (SBD) speed estimates by deployment, for each deployment with at least 10 movement sequences. n = number of sequences retained after quality filtering; SBD speed = harmonic mean speed corrected for size-biased sampling (Rowcliffe et al. 2016); SE = standard error; 95% CI = bootstrap confidence interval (2.5th and 97.5th percentiles, 1,000 replicates).

| Deployment | Species | n | SBD speed (m/s) | SE | 95% CI (m/s) |
|---|---|---|---|---|---|
| Z18 | Mule Deer | 14 | 1.324 | 1.098 | 0.870–2.463 |
| Z20 | Mule Deer | 12 | 0.533 | 0.466 | 0.356–1.020 |
| H25 | Red Squirrel | 10 | 0.040 | 0.075 | 0.014–0.454 |
| Y17 | Red Squirrel | 19 | 0.380 | 0.096 | 0.259–0.580 |
| Y29 | Red Squirrel | 28 | 0.095 | 0.098 | 0.038–0.500 |
| D26 | Snowshoe Hare | 10 | 0.368 | 0.167 | 0.246–0.633 |

| Deployment | Species | n | SBD speed (m/s) | SE | 95% CI (m/s) |
|---|---|---|---|---|---|
| C29 | White-tailed Deer | 15 | 0.693 | 0.264 | 0.527–0.998 |
| E16 | White-tailed Deer | 22 | 0.158 | 0.079 | 0.096–0.301 |
| E18 | White-tailed Deer | 13 | 0.505 | 0.098 | 0.354–0.751 |
| E19 | White-tailed Deer | 18 | 0.318 | 0.163 | 0.202–0.517 |
| E21 | White-tailed Deer | 24 | 0.073 | 0.155 | 0.037–0.207 |
| E28 | White-tailed Deer | 10 | 0.390 | 0.083 | 0.286–0.580 |
| E94 | White-tailed Deer | 12 | 0.414 | 0.273 | 0.264–0.709 |
| E96 | White-tailed Deer | 17 | 0.073 | 0.052 | 0.042–0.203 |
| F13 | White-tailed Deer | 17 | 0.118 | 0.076 | 0.069–0.289 |
| F18 | White-tailed Deer | 22 | 0.146 | 0.061 | 0.100–0.242 |
| F20 | White-tailed Deer | 16 | 0.049 | 0.113 | 0.030–0.108 |
| F22 | White-tailed Deer | 20 | 0.172 | 0.105 | 0.111–0.301 |
| F24 | White-tailed Deer | 23 | 0.240 | 0.160 | 0.171–0.372 |
| F26 | White-tailed Deer | 15 | 0.492 | 0.154 | 0.336–0.788 |
| F4 | White-tailed Deer | 19 | 0.053 | 0.065 | 0.029–0.115 |
| F5 | White-tailed Deer | 19 | 0.197 | 0.270 | 0.135–0.306 |
| F8 | White-tailed Deer | 35 | 0.157 | 0.081 | 0.103–0.280 |
| F9 | White-tailed Deer | 22 | 0.174 | 0.094 | 0.122–0.277 |
| G11 | White-tailed Deer | 39 | 0.139 | 0.097 | 0.105–0.203 |
| G12 | White-tailed Deer | 19 | 0.532 | 0.098 | 0.360–0.843 |
| G14 | White-tailed Deer | 21 | 0.643 | 0.225 | 0.464–0.964 |
| G2 | White-tailed Deer | 39 | 0.046 | 0.077 | 0.032–0.073 |
| G4 | White-tailed Deer | 60 | 0.308 | 0.089 | 0.210–0.485 |
| G6 | White-tailed Deer | 16 | 0.124 | 0.152 | 0.056–0.423 |
| H4 | White-tailed Deer | 15 | 0.186 | 0.317 | 0.089–0.603 |
| H5 | White-tailed Deer | 11 | 0.165 | 0.043 | 0.106–0.278 |
| I23 | White-tailed Deer | 11 | 0.272 | 0.081 | 0.192–0.422 |
| I24 | White-tailed Deer | 13 | 0.235 | 0.092 | 0.169–0.350 |
| I28 | White-tailed Deer | 16 | 0.158 | 0.064 | 0.094–0.304 |
| I30 | White-tailed Deer | 51 | 0.175 | 0.077 | 0.137–0.232 |
| Z18 | White-tailed Deer | 16 | 1.252 | 0.907 | 0.845–2.012 |
| Z20 | White-tailed Deer | 10 | 0.403 | 0.334 | 0.228–0.891 |

**Table S3.** DPT scene depth by deployment, defined as the mean depth value estimated by the Dense Prediction Transformer across the full scene at each deployment, extracted independently of any animal detections, serving as a proxy for habitat openness. See data availability statement for table.

**Table S4.** Effective detection distance (EDD) estimates by species from the joint multi-species hierarchical model, ordered by number of detections.

| Species | Deployments | Detections | Mean EDD (m) | SD | Range (m) |
|---|---|---|---|---|---|
| White-tailed deer | 138 | 2245 | 6.18 | 3.03 | 0.78–13.9 |
| American black bear | 99 | 240 | 4.91 | 2.72 | 0.64–14.5 |
| Red squirrel | 35 | 217 | 4.07 | 2.38 | 0.58–9.2 |
| Mule deer | 64 | 208 | 6.68 | 3.15 | 0.77–15.8 |
| Snowshoe hare | 48 | 176 | 4.47 | 2.46 | 1.01–11.6 |
| Elk | 41 | 126 | 7.08 | 3.12 | 1.67–15.9 |

**Table S5.** Species-specific coordinate anchoring thresholds and selection method (see Section S2).

| Body size class | Species | Threshold (m) | Method |
|---|---|---|---|
| Large ungulate | White-tailed deer | 0.75 | Override from 0.3 |
| Large ungulate | Elk | 1.0 | Kneedle |
| Large ungulate | Mule deer | 0.5 | Override from 0.3 |
| Large ungulate | Moose | 0.5 | Body-size default |
| Medium carnivore/omnivore | American black bear | 0.75 | Kneedle |
| Medium carnivore/omnivore | Bobcat | 1.0 | Kneedle |
| Medium carnivore/omnivore | Puma | 0.3 | Body-size default |

| Body size class | Species | Threshold (m) | Method |
|---|---|---|---|
| Medium carnivore/omnivore | Coyote | 0.3 | Body-size default |
| Small mammal | Red squirrel | 0.1 | Kneedle |
| Small mammal | Snowshoe hare | 0.1 | Body-size default |
| Bird | Wild turkey | 0.5 | Kneedle |

##### Figures

**Figure S1.** Distance-dependent depth estimation precision. Each point represents one multi-frame sequence; the LOESS smooth shows increasing coordinate jitter with distance.

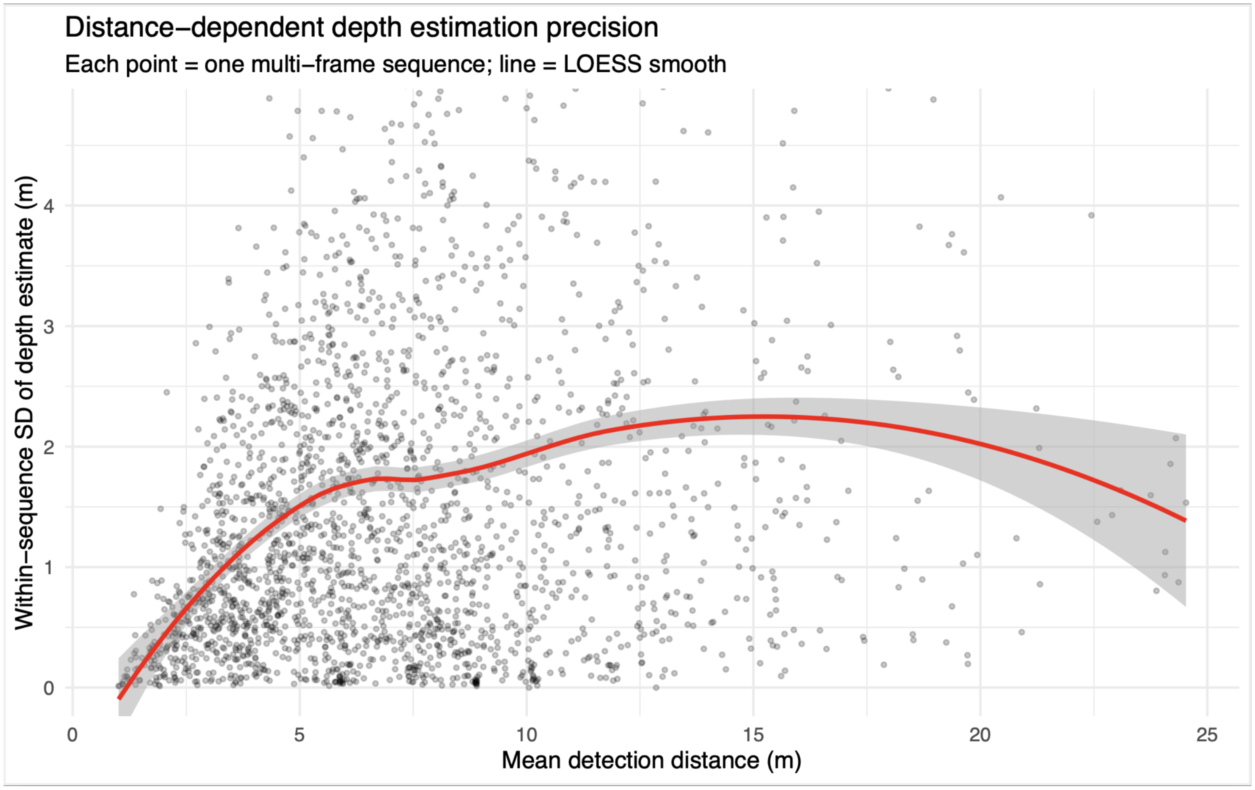


**Figure S2.** Tortuous path filtering example. Left: filtered sequence with extreme tortuosity from AI coordinate noise. Right: retained sequence with biologically plausible path.

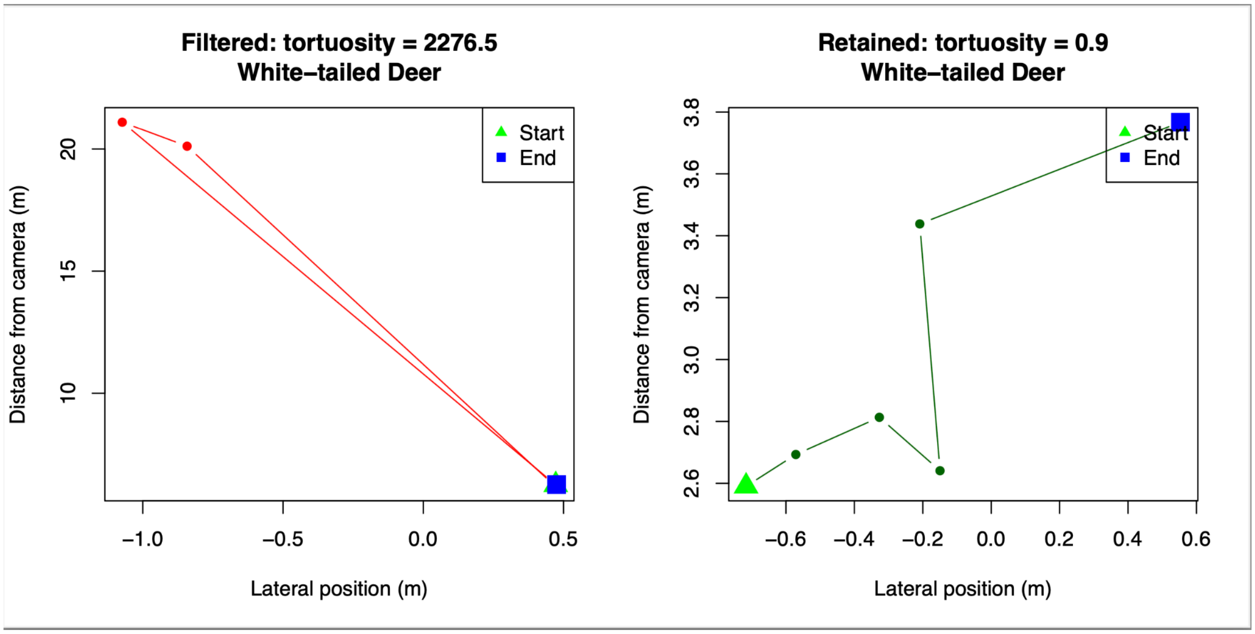


**Figure S3.** Empirical anchoring threshold selection via Kneedle elbow detection. Size-biased distribution (SBD) speed estimates (m/s, top) and number of retained sequences (bottom) across eight candidate anchoring thresholds for white-tailed deer and American black bear. The vertical line indicates the final selected threshold. Where the Kneedle algorithm and final threshold

differ, the algorithmic

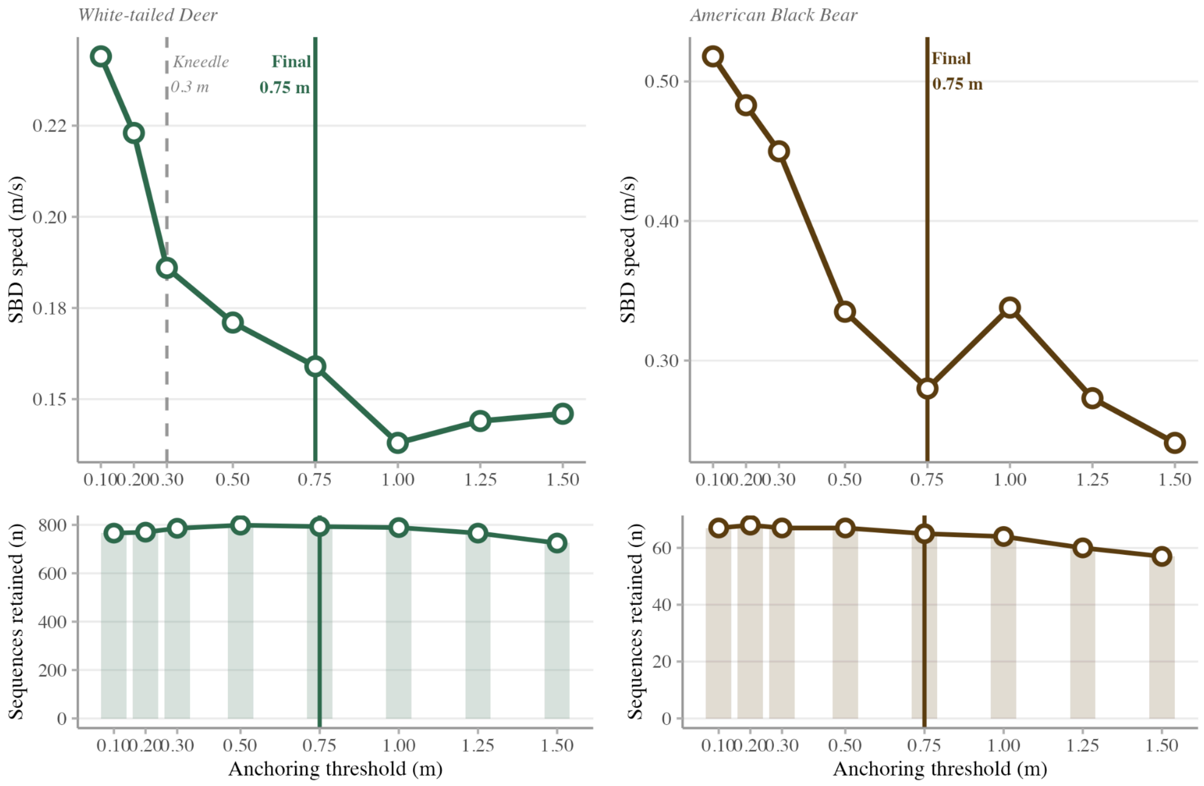


**Figure S4.** Effect of coordinate anchoring on movement speed distributions for American black bear and white-tailed deer. Curves show kernel density estimates of sequence-level speeds derived with (orange) and without (blue) species-specific anchoring thresholds. Anchoring reduces the influence of sub-threshold displacements treated as coordinate noise, shifting the distribution leftward and dampening right-tail inflation.

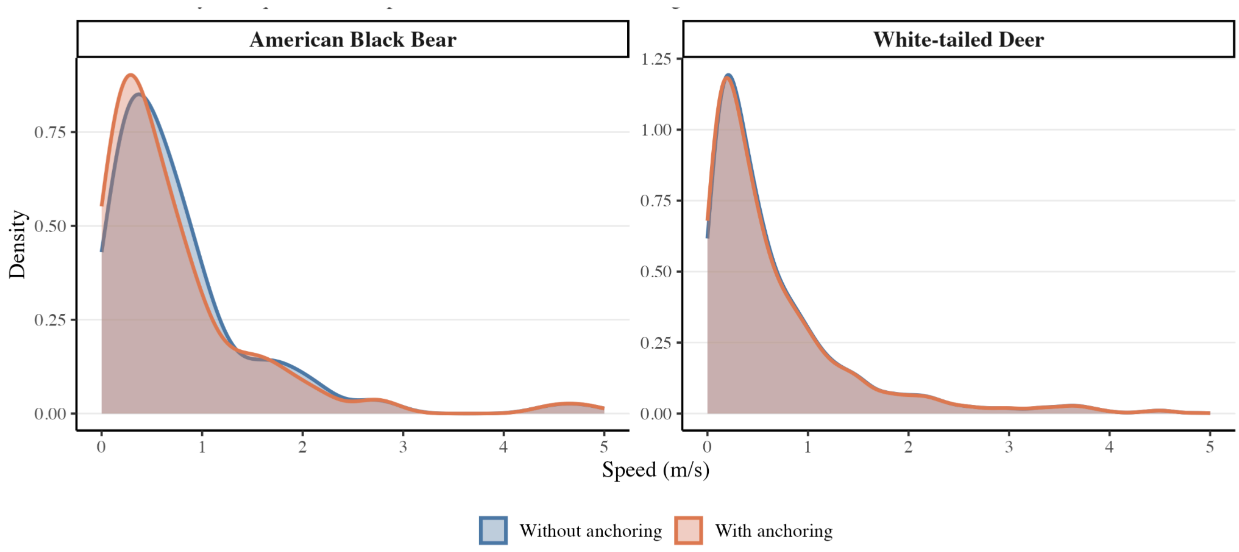


**Figure S5.** Calibration reference images illustrating the calibration setup for a single deployment, with a high-contrast distance sign held at known distances along the camera's viewing axis.

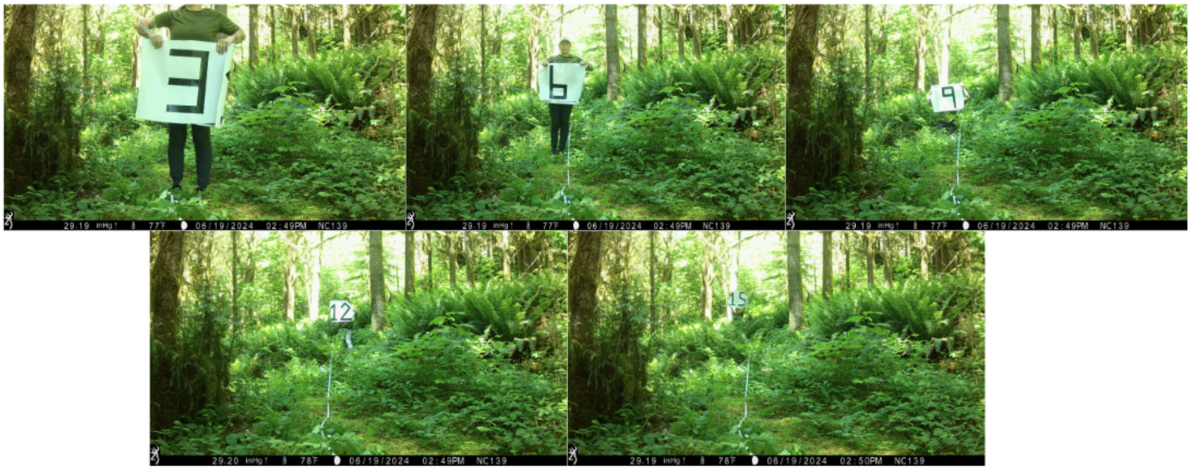

Distributed Computing Systems Workshops:166–171.

https://doi.org/10.1109/ICDCSW.2011.20.
